# Within-Subject Optogenetic Model Reveals Spatiotemporal Cortical Reorganization in Artificial Vision

**DOI:** 10.64898/2026.08.19.745677

**Authors:** Xurong Gao, Wenjie Ren, Baisong Zhang, Yueqian Zhou, Guangda Hu, Yuchen Xu, Chengpeng Chai, Chuanqing Wang, Yong Zou, Lifeng Wang, Yun-Hsuan Chen, Jie Yang, Mohamad Sawan

**Affiliations:** CenBRAIN Neurotech, Westlake University, 600 Dunyu Road, Xihu District, Hangzhou, Zhejiang, 310030, China; Beijing Institute of Radiation Medicine; Beijing, 100850, China

**Keywords:** Artificial vision, Visual prosthesis, Neural representation, Retinal optogenetics, Deep learning, Primary visual cortex

## Abstract

Visual prostheses can elicit simple percepts such as letter forms through dynamically patterned stimulation, yet naturalistic dynamic vision remains out of reach despite hardware fully capable of high-resolution temporal control. This persistent gap suggests the bottleneck lies in how artificial input is represented by the cortex itself. A fundamental question remains unresolved: when artificial vision bypasses natural retinal encoding, what is lost in cortical representation? We address this question in this paper using a retinal optogenetic mouse model that isolates the consequence of bypassing retinal encoding while preserving downstream pathways. Using within-subject V1 cortex electrophysiology and CNN-based decoding under identical dynamic stimuli, we reveal dissociable, dimension-specific gaps. Temporally, artificial vision bypassed the frequency-dependent compression imposed by natural vision, extending stimulus-locked entrainment bandwidth. Spatially, direction-selective representations underwent systematic remapping, with disrupted phase organization and reduced encoding regularity. These findings show that bypassing natural retinal encoding reorganizes cortical representation along dissociable spatiotemporal dimensions, establishing a within-subject framework that renders such structural divergences easy to locate and quantifiable.

## Introduction

By exploiting stable phosphene topography to design temporally patterned sequences ^1,2^, visual cortical prostheses can now reliably elicit simple percepts, such as letter forms, in both sighted and blind subjects ^3,4^. Building on this success, a prevailing assumption has emerged across the field: scaling these temporal sequences in resolution and complexity should naturally extend to continuous, naturalistic dynamic vision ^5–7^. However, although current implanted visual prostheses have already reached stimulation capabilities sufficient for dynamic visual scenarios at the hardware level, they have yet to achieve continuous dynamic percepts ^8–11^. This discrepancy raises a key question: how are temporally encoded artificial stimuli represented and integrated in the cortex, and why do they fail to consistently give rise to robust dynamic percepts ^4^? One possibility is that the cortex simply fails to track the temporal structure of the artificial visual input with sufficient fidelity. Another is that bypassing natural encoding confronts the cortex with an incompatible signal format, forcing a structurally altered representation that distorts the resulting percept ^12^.

To systematically test these hypotheses, we established a retinal optogenetic model in mice as a representative system for artificial visual prosthetics (Fig. 1a,b) ^13–15^. We performed longitudinal electrophysiological recordings in the primary visual cortex (V1), enabling a direct, within-subject comparison of neural representations across two physiological stages (Fig. 1a). During the initial baseline stage with intact vision, we mapped V1 responses to fundamental temporal and spatial dynamic stimuli (Fig. 1c,d) ^16,17^. Subsequently, following induced blindness and optogenetic intervention, we delivered mathematically equivalent patterns of optical stimulation (Fig. 1b–d) ^18,19^. By contrasting these intact and artificial visual states within the same individuals, we aimed to pinpoint the exact divergence in spatiotemporal representations. To capture these complex multidimensional dynamics, we integrated classical electrophysiology with a One-Dimensional Convolutional Neural Network (1D-CNN) ^20–22^, enabling robust decoding of the underlying spatiotemporal features from cortical responses (Fig. 1e,f) ^23^.

**Fig. 1.**
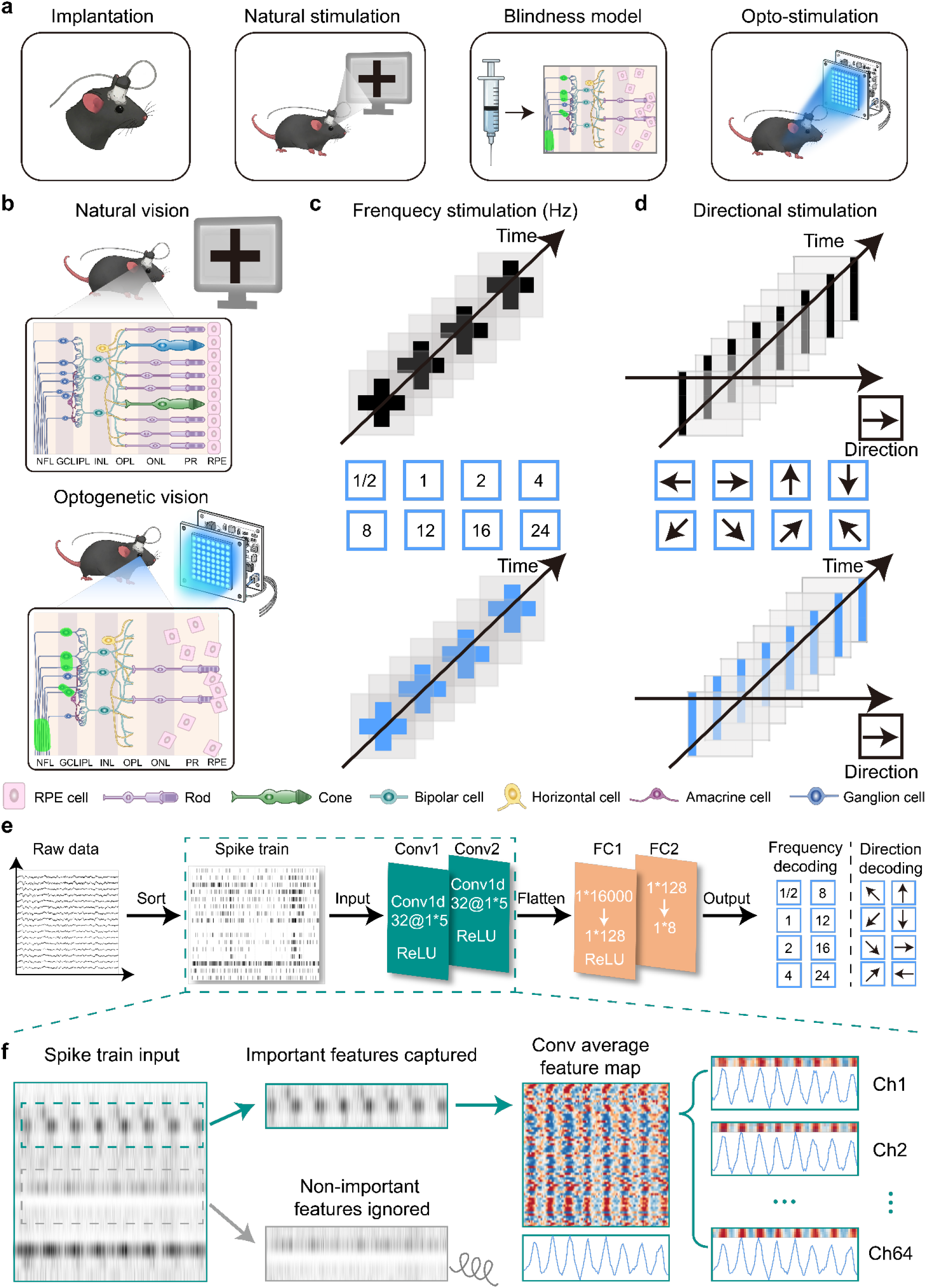
Experimental framework for within-subject comparison of natural and optogenetic artificial vision. **a,** Schematic of the longitudinal experimental design. Mice were implanted with cortical recording electrodes and first recorded under intact normal vision during screen-based visual stimulation. A blindness model was then induced, and an Adeno-Associate Virus (AAV) vector was delivered to confer photosensitivity to retinal ganglion cells. The same mice were subsequently tested under optogenetic visual stimulation. Opto denotes optogenetic stimulation. **b,** Comparison of the two visual input pathways. Under natural vision, screen-presented stimuli are processed through the intact retinal circuitry before reaching V1. Under optogenetic artificial vision, photoreceptors are disrupted and retinal ganglion cells become photosensitive, allowing patterned light stimulation to evoke ganglion-cell activity that is transmitted to the visual cortex. **c,** Temporal-rate stimulation paradigm. Visual sequences presented under natural vision were converted into equivalent optogenetic stimulation patterns. The Steady State Visually Evoked Potential (SSVEP) flicker frequencies were 0.5, 1, 2, 4, 8, 12, 16 and 24 Hz. **d,** Directional stimulation paradigm. Moving visual sequences presented under natural vision were converted into equivalent patterned optogenetic stimulation sequences. The scanning-grating directions were 0°, 45°, 90°, 135°, 180°, 225°, 270° and 315°. **e,** Electrophysiology and decoding pipeline. Raw V1 recordings were spike-sorted into population spike trains and used as input to a one-dimensional convolutional neural network for frequency and direction decoding. **f,** Schematic of 1D-CNN-based feature extraction. Convolutional filters captured stimulus-relevant temporal structure from population spike trains while suppressing less informative components, generating feature maps used to analyze cortical encoding of temporal and spatial stimulus structure.

Here, we report a distinct spatiotemporal dissociation in the cortical processing of artificial vision ^24,25^. Our analysis demonstrates that the difficulty in synthesizing robust dynamic percepts does not arise from a temporal tracking deficit, but rather stems from an absence of coordinated spatial phase organization ^26^. Contrary to the assumption that artificial inputs suffer from reduced transmission fidelity, our neural manifold analysis and CNN feature mapping reveal that the optogenetic pathway actually resolves temporal stimulation sequences more discretely than the intact visual system ^27,28^. Specifically, normal vision appears to utilize an efficient coding strategy, allocating high discriminability to low-frequency signals while compressing ascending mid-to-high frequency stimuli into grouped, overlapping state spaces. The artificial pathway bypasses this natural compression; it distributes representational resources evenly across the tested frequency spectrum, maintaining distinct boundary separations and preserving high-frequency tracking where natural encoding typically exhibits information loss ^23^.

Crucially, however, we observed that this uncompressed temporal capacity is inherently decoupled from functional spatial encoding. During natural spatial sequence processing, the visual cortex employs highly unified, phase-differentiated neural representations to continuously encode spatial information. Conversely, under equivalent artificial visual stimuli, the cortical spatial representation undergoes a profound structural reorganization. Electrophysiological analysis reveals that while direction selectivity persists, it undergoes a systematic orthogonal remapping that fundamentally shifts the population-level tuning geometry ^29^. Furthermore, deep learning-based feature decoding demonstrates that this remapped spatial representation loses its intrinsic phase organization. This deficit is characterized by the disruption of stable temporal response structures, rendering the encoded information highly disorganized and significantly less separable. Taken together, this analysis demonstrates that bypassing natural encoding creates a systematic mismatch with the inherent spatiotemporal decoding architecture of the visual cortex ^12^. This mechanistic mismatch clarifies a critical gap in achieving robust dynamic artificial vision. Ultimately, the within-subject comparative framework established here offers a practical diagnostic paradigm for evaluating and refining the spatiotemporal encoding strategies of future visual prostheses.

## Results

We established a within-subject, two-stage experimental paradigm to directly compare cortical representations of natural and artificial vision (Supplementary Fig. S1). Successful blinding and optogenetic restoration were jointly validated through V1 electrophysiological responses, decoding performance, and retinal histology (Supplementary Fig. S2). The dynamic properties of optogenetic drive in V1 were further characterized across light intensities, frequencies, and pulse durations (Supplementary Fig. S3). Building upon this functionally validated foundation, we next investigated the underlying spatiotemporal encoding architectures in V1, studying how population-level feature representations structurally diverge when visual input bypasses natural retinal processing.

### Artificial vision relieves frequency-dependent compression of V1 population states

To determine whether artificial vision degrades cortical representations of stimulus dynamics, we first examined how V1 population activity encoded the temporal rate of visual sequences under natural and optogenetically driven visual states. In the same cohort of mice (n = 5), drifting grating stimuli were presented at eight temporal update rates (2, 4, 8, 12, 16, 20, 24, 28Hz) during the intact-vision baseline and again after optogenetic intervention using matched artificial stimulation patterns. Trial-wise V1 spike-train responses were projected into a two-dimensional embedding space using t-Distributed Stochastic Neighbor Embedding (t-SNE) to visualize the geometry of population responses across temporal rates ^30^.

Under normal vision, V1 population states exhibited progressive frequency-dependent compression (Fig. 2a,c). In four mice (#1, #3, #4 and #5), slow update rates (2–8 Hz) formed relatively discrete clusters, whereas responses became increasingly adjacent or overlapping at intermediate rates (12–16 Hz) and converged strongly into shared state spaces at the fastest rates (20–28 Hz) ^31^. Mouse #2 showed weaker low-frequency separation, but still followed the same trend of increasing overlap with stimulus rate. Consistent with an efficient coding strategy, retina-mediated vision preserved highly separable representations for slow temporal sequences while compressing faster visual dynamics into grouped representational resources.

**Fig. 2.**
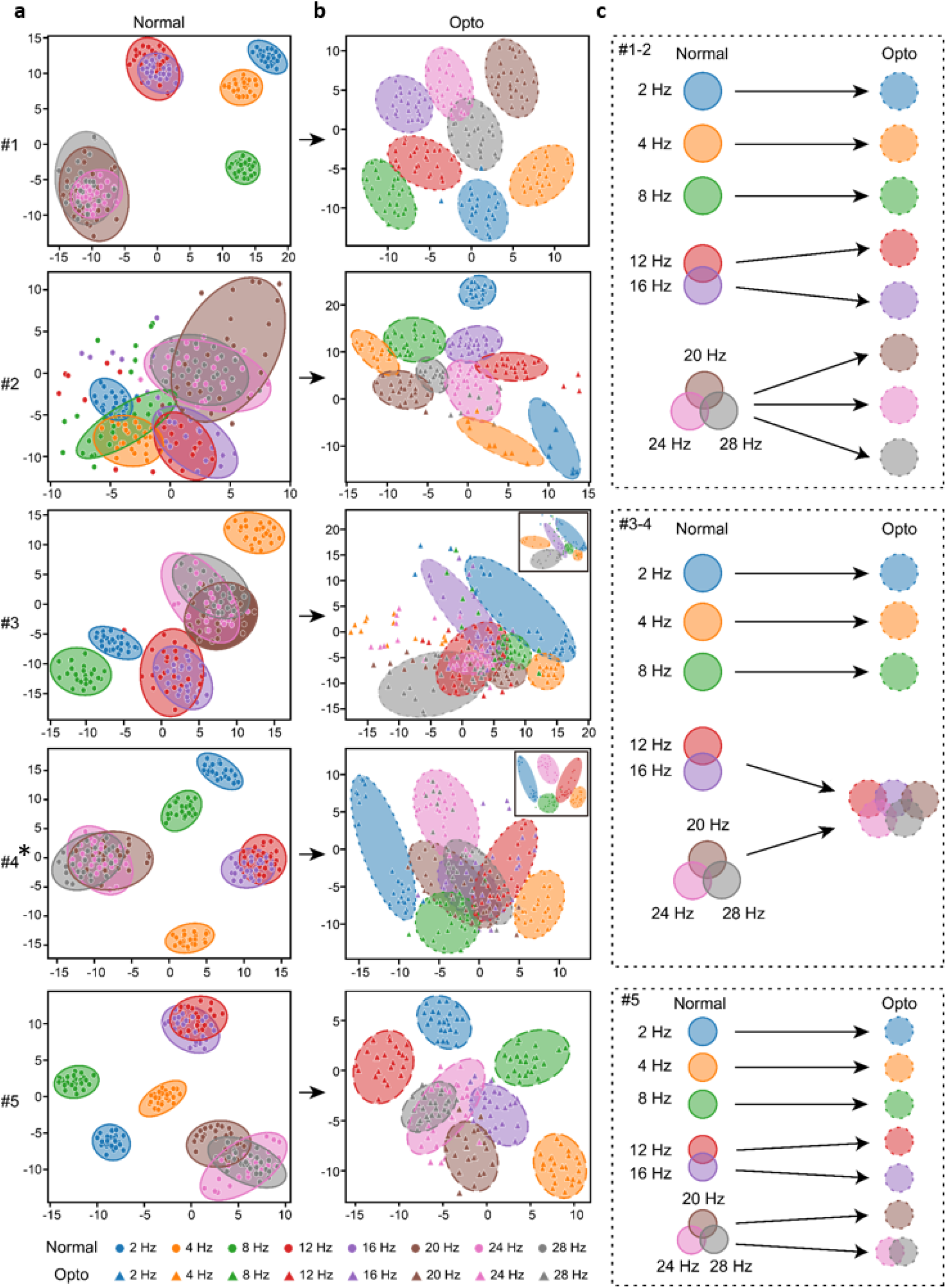
t-SNE visualization of V1 population states across temporal update rates under natural and optogenetic vision. **a,** Trial-wise t-SNE embeddings of V1 population spike-train responses under natural vision in five mice. Each point represents one trial. Colors indicate temporal update rates of drifting grating sequences (2 to 28 Hz). Circular markers denote trials recorded under natural vision, and ellipses outline the trial distribution for each frequency condition. The clustering architecture reveals a progressive frequency-dependent compression at higher temporal rates. The axes represent the first two dimensions of the t-SNE embedding to visualize relative population-state geometry. **b,** Trial-wise t-SNE embeddings of V1 population responses from the same mice under optogenetic artificial vision. Triangular markers denote trials recorded under optogenetic stimulation. Colors indicate the same temporal update rates as in a. The embeddings demonstrate that artificial vision fundamentally reorganizes the population geometry. It either bypasses natural compression to distinctively isolate high-frequency domains (mice #1, #2, and #5) or results in extensive structural smearing (mice #3 and #4). **c,** Schematic summary of the frequency-state organization. Solid circles on the left represent natural vision states, illustrating graded overlap and representational compression. Dashed circles on the right illustrate the corresponding optogenetic states. Arrows highlight the structural uncoupling of frequency domains, showing how artificial drive segregates previously compressed high-frequency conditions in the majority of subjects. The asterisk on mouse #4 denotes confirmed low AAV expression (Supplementary Fig. S4), which likely accounts for the technical variation and disorganized overlap observed in this subject.

Optogenetic artificial vision fundamentally disrupted this natural population-state geometry. In the majority of the cohort (mice #1, #2, and #5), optogenetic stimulation distinctly isolated the high-frequency conditions that were previously compressed under natural vision. These animals formed highly separable domains across the tested spectrum, with residual overlap in mouse #5 strictly confined to the two fastest rates. In contrast, mice #3 and #4 exhibited structural smearing and increased trial-by-trial variance at higher frequencies. This likely represents reasonable variation resulting from technical limitations, such as insufficient viral transfection, or inherent subject-specific heterogeneity. Despite these expected individual differences, the dominant trend demonstrates that effective artificial drive bypasses natural frequency-dependent compression, enforcing artificially distinct boundaries between temporal states in V1.

### Artificial vision extends the temporal bandwidth of cortical entrainment

In Section A, we observed that artificial vision uncompresses high-frequency states. To test whether this reflects a genuine capacity for rapid temporal tracking, we evaluated V1 responses to pure temporal flickers (SSVEP) ^31^. This stimulus isolates temporal processing by removing the spatial movement inherent to drifting gratings. CNN-derived feature maps (Fig. 3a) showed that both modalities tracked low frequencies reliably. At higher frequency bands, however, natural vision progressively lost its temporal tuning, whereas artificial vision retained partial rhythmic tracking. Consistent patterns were observed across the remaining subjects (Supplementary Fig. S6).

**Fig. 3.**
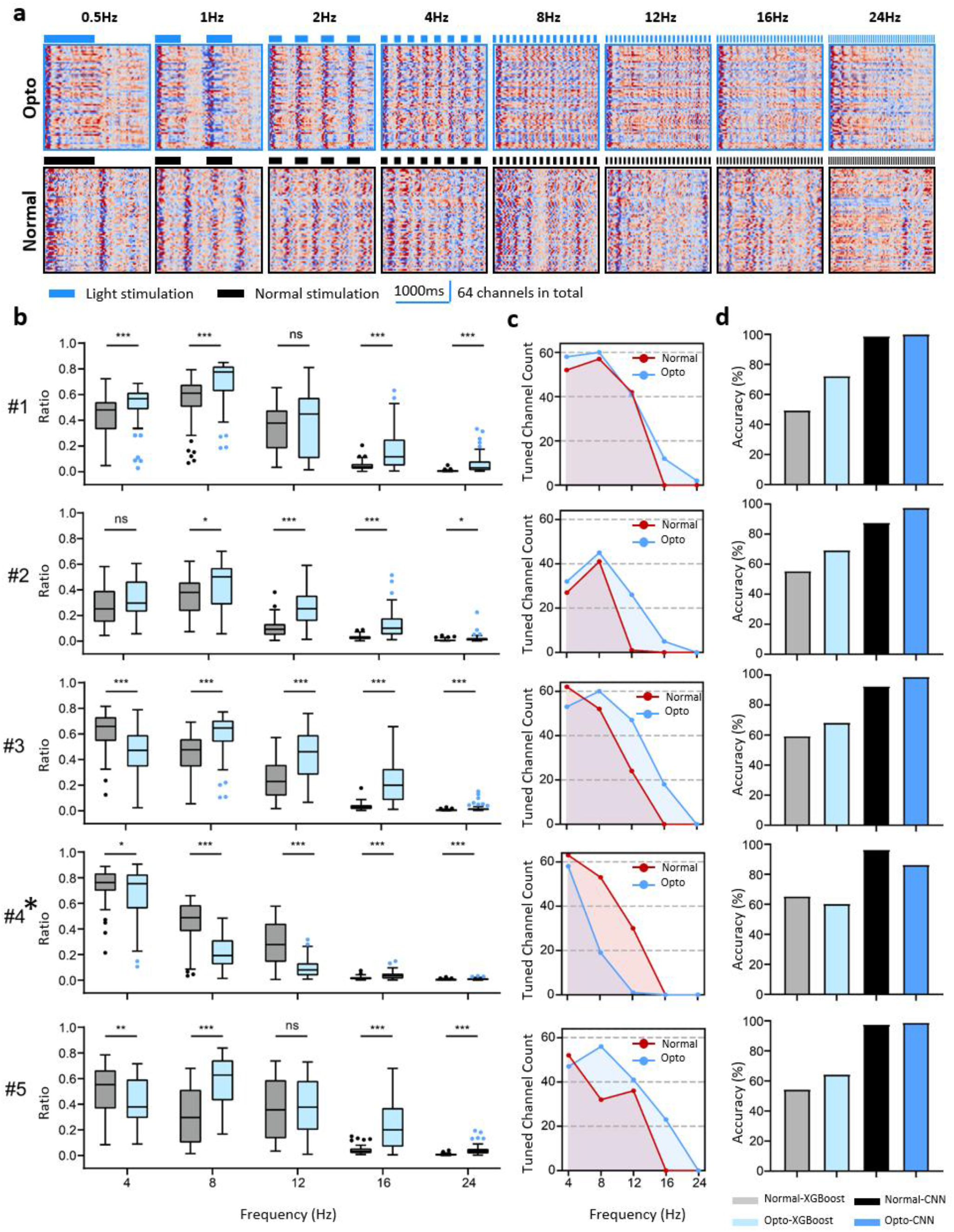
Artificial vision extends the temporal bandwidth of cortical entrainment. **a,** Representative CNN feature maps extracted from V1 responses to SSVEP flicker stimulation under optogenetic artificial vision and natural vision. Columns show flicker frequencies of 0.5, 1, 2, 4, 8, 12, 16 and 24 Hz. Each heat map shows the second convolutional-layer output of the two-layer 1D-CNN, consisting of 64 model-derived feature channels. Blue and black bars indicate the on/off periods of optogenetic light stimulation and natural visual stimulation, respectively. Scale bar, 1,000 ms. **b,** Stimulus-frequency power in CNN-derived feature channels for five mice. For each model output feature channel and frequency condition, the ratio of spectral power at the stimulus-frequency band to total spectral power was calculated after Fourier transformation. Each point represents one model-derived feature channel. Box plots show channel-wise distributions. ns, not significant; *p < 0.05, **p < 0.01, and ***p < 0.001. **c,** Number of frequency-tuned CNN feature channels across flicker frequencies in each mouse. Tuned channels were defined as model output feature channels in which the stimulus-frequency component accounted for more than 30% of total spectral power. Red, natural vision; blue, optogenetic stimulation. The maximum channel count is 64, determined by the second convolutional-layer output. **d,** Decoding accuracy for SSVEP flicker-frequency classification from V1 population responses using XGBoost and CNN classifiers under natural and optogenetic visual conditions. The legend indicates the classifier type and visual condition for each bar. The asterisk on #4 (#4*) denotes low AAV expression in this subject (Supplementary Fig. S4), which likely accounts for its reduced optogenetic performance.

Spectral analysis quantified this extended tracking capability (Fig. 3b). The frequency-specific power decayed rapidly under natural vision as frequencies increased, whereas it remained strongly preserved under artificial drive. Similarly, at the population level, the number of V1 channels maintaining strong temporal tuning dropped toward floor levels under natural vision but was largely sustained across most frequency bands under artificial vision (Fig. 3c). The protocol for characterizing the neural representations of CNN feature maps under SSVEP stimulation is shown in (Supplementary Fig. S5). Because artificial vision preserves these distinct temporal features across a wider spectrum, stimulus-frequency decoding achieved significantly higher accuracy in the mid-to-high ranges compared to natural vision, where feature blurring degraded discriminability (Fig. 3d). The sole exception was mouse #4, where confirmed low opsin expression simply constrained the local drive, representing an expected technical variance (Supplementary Fig. S4).

Crucially, combining the results from Sections A and B directly addresses the hypothesis raised in our introduction. These findings demonstrate that the cortex does not simply fail to track the temporal structure of the artificial visual input with sufficient fidelity. On the contrary, because artificial vision bypasses the retinal low-pass filter, its temporal fidelity is actually enhanced, granting the cortex a higher frequency ceiling. Although the brain’s downstream networks may not readily adapt to these uncompressed signal formats lacking the equivalent processing of natural visual circuits, this discovery points to a clear optimization strategy: artificially simulating the natural compression mechanism to format the prosthetic input. Furthermore, having ruled out a deficit in temporal fidelity as the primary bottleneck, we next sought to investigate whether the distortion of dynamic perception emerges instead within the spatial domain.

### Artificial vision induces a systematic orthogonal remapping of spatial tuning geometry

Having established that artificial vision preserves temporal fidelity, we next asked whether it also preserves the spatial structure of motion coding. We presented drifting gratings moving in eight directions and recorded direction-selective V1 responses under natural and optogenetically driven visual states. In a representative animal, trial-averaged responses across 16 recording channels showed clear direction tuning under natural vision, with identifiable preferred directions and graded response modulation across the stimulus set (Fig. 4a). Optogenetic stimulation also elicited direction-selective responses, indicating that artificial vision did not simply abolish motion-direction tuning. However, the preferred directions and tuning profiles of individual channels differed substantially between the two visual states. Directions that strongly activated a channel under natural vision often evoked weaker or suppressed responses under optogenetic stimulation, whereas previously weaker directions could become dominant. Thus, optogenetic drive preserved direction selectivity but altered the mapping between stimulus direction and cortical response.

**Fig. 4.**
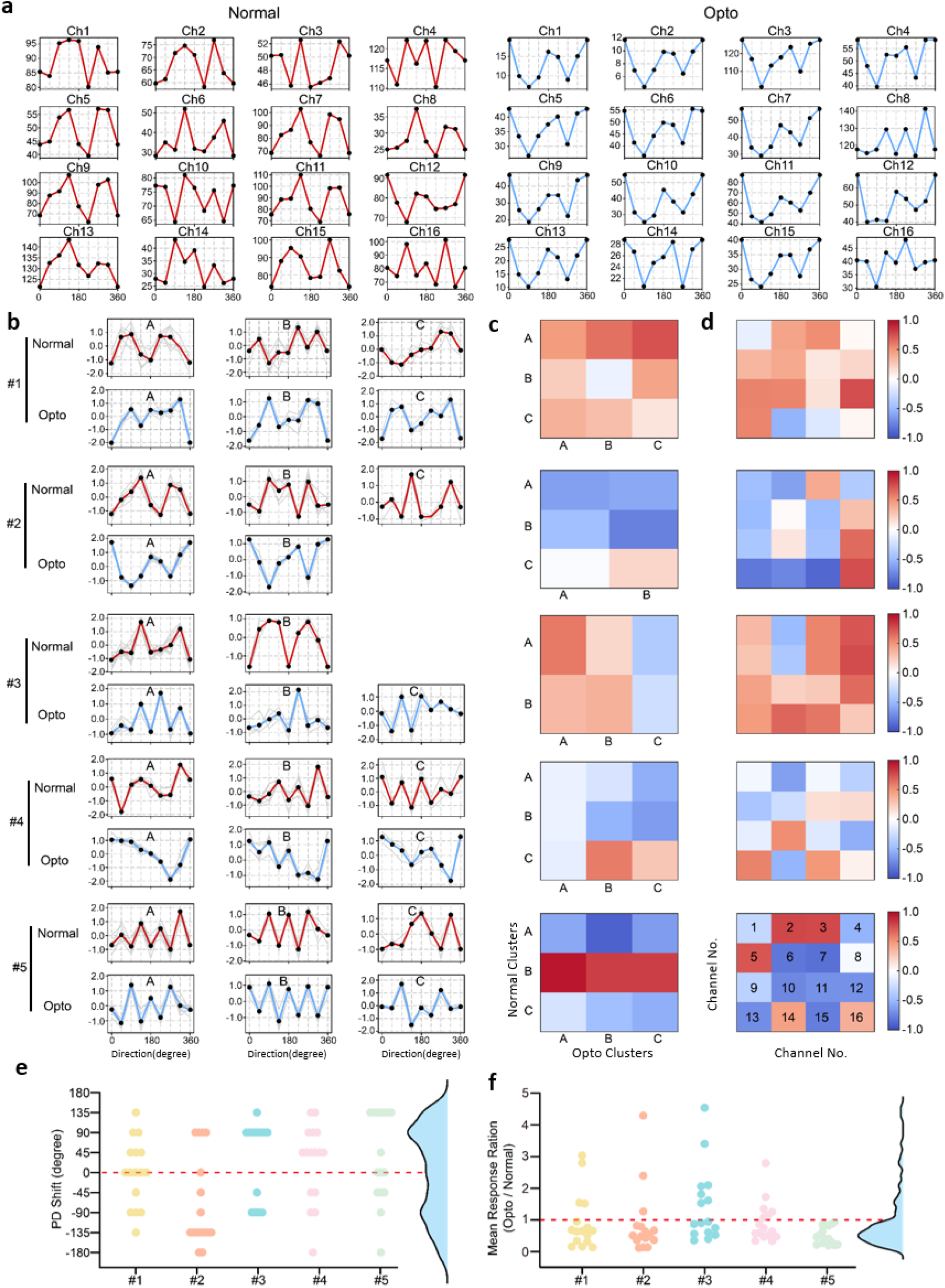
Artificial vision induces orthogonal remapping of direction selectivity in V1. **a,** Example direction tuning curves from 16 recording channels in one representative mouse under natural vision and optogenetic artificial vision. Drifting gratings were presented in eight directions from 0° to 315° in 45° steps, with 50 trials per direction. Red, natural vision; blue, optogenetic stimulation. Opto denotes optogenetic stimulation. **b,** Cluster-averaged direction tuning profiles for five mice. Individual channel tuning curves were grouped by unsupervised clustering separately under natural and optogenetic conditions. Gray traces show individual channel profiles; colored traces show cluster means. **c,** Pearson correlation matrices between natural-vision and optogenetic cluster-averaged tuning curves. Rows indicate natural-vision clusters, and columns indicate optogenetic clusters. Warm and cool colors indicate positive and negative correlations, respectively. **d,** Channel-wise Pearson correlations between natural-vision and optogenetic direction tuning curves, arranged according to the 4 x 4 recording-channel layout. The lower-right schematic indicates channel numbering. **e,** Preferred direction shift (PD shift) for individual channels between visual conditions. Each point represents one recording channel; values near 0° indicate preserved preferred direction, whereas values near +/-90° indicate an approximately orthogonal shift. The marginal density plot summarizes the distribution. **f,** Mean response ratio between optogenetic and natural visual conditions for individual channels. Ratios were calculated from mean firing rates averaged across all eight directions and trials. The dashed line marks a ratio of 1, and the marginal density plot summarizes the distribution.

We next examined whether this remapping extended beyond individual channels to the population structure of direction coding. Unsupervised clustering of direction tuning profiles revealed well-defined direction-selective ensembles under natural vision, with cluster-averaged tuning curves distributed across the tested directions (Fig. 4b). Under optogenetic stimulation, direction-selective clusters were still present, but their mean tuning profiles were markedly reorganized relative to natural vision. Pairwise correlations between cluster-averaged tuning curves further showed that the relationships among direction-selective groups changed across visual states, with altered similarity and orthogonality between clusters (Fig. 4c). Consistently, channel-wise correlations between natural and optogenetic tuning curves were low or negative for most recording sites, indicating that individual channels generally did not preserve their original direction preferences under artificial vision (Fig. 4d).

To determine whether this reorganization reflected random instability or a structured transformation, we quantified the shift in preferred direction for each channel across visual states (Fig. 4e). If direction preferences were preserved, the distribution of shifts would be expected to cluster near 0°. Instead, shifts were concentrated near ±90°, forming a bimodal distribution. This indicates that optogenetic stimulation did not merely introduce nonspecific variability or uniformly degrade direction coding. Rather, it systematically remapped preferred directions toward orthogonal motion axes. Preservation of the original direction preference was rare, suggesting that artificial vision imposes a distinct spatial coding format on V1 motion representations.

Finally, we examined whether direction remapping was accompanied by changes in response magnitude (Fig. 4f). For each channel, we calculated the ratio of its mean response under optogenetic stimulation to that under natural vision. The resulting distribution was not consistent with a simple global scaling of cortical activity. Although many channels showed ratios below unity, indicating reduced average responses under optogenetic drive, a subset of channels showed ratios above unity, reflecting selective enhancement. Thus, artificial vision reshaped the gain landscape of V1 responses in a channel-specific manner. Together with the orthogonal shifts in preferred direction, this indicates that optogenetic stimulation does not merely weaken or amplify natural direction coding. Instead, it reformats V1 motion representations by jointly remapping direction preference and redistributing response gain.

### Artificial vision disrupts the intrinsic phase organization of spatial encoding, degrading decodability

The preceding analyses showed that artificial vision preserves direction-selective responses in V1 but reorganizes their spatial mapping. We next asked whether these altered responses still formed reliable, decodable representations of motion direction. To this end, we trained 1D-CNN decoders on V1 activity and examined the learned feature representations under natural and optogenetically driven visual states.

CNN feature maps revealed a clear difference in feature organization between the two conditions. Under natural vision, drifting gratings evoked structured feature patterns across motion directions, with dense activation within the post-stimulus response window and a reproducible double-peak temporal profile in many channels (Fig. 5a, b). The timing and shape of these feature responses varied systematically across directions, indicating that motion direction was represented by organized temporal patterns. Under optogenetic stimulation, direction-related activity was still present, but the feature maps were less regular. Activation was sparser and less temporally aligned, and the double-peak structure was weaker or absent in many channels (Fig. 5a, c). Consistent patterns were observed across the remaining subjects (Supplementary Fig. S7). Thus, artificial vision did not eliminate motion-related activity, but it reduced the temporal organization of the learned direction features.

**Fig. 5.**
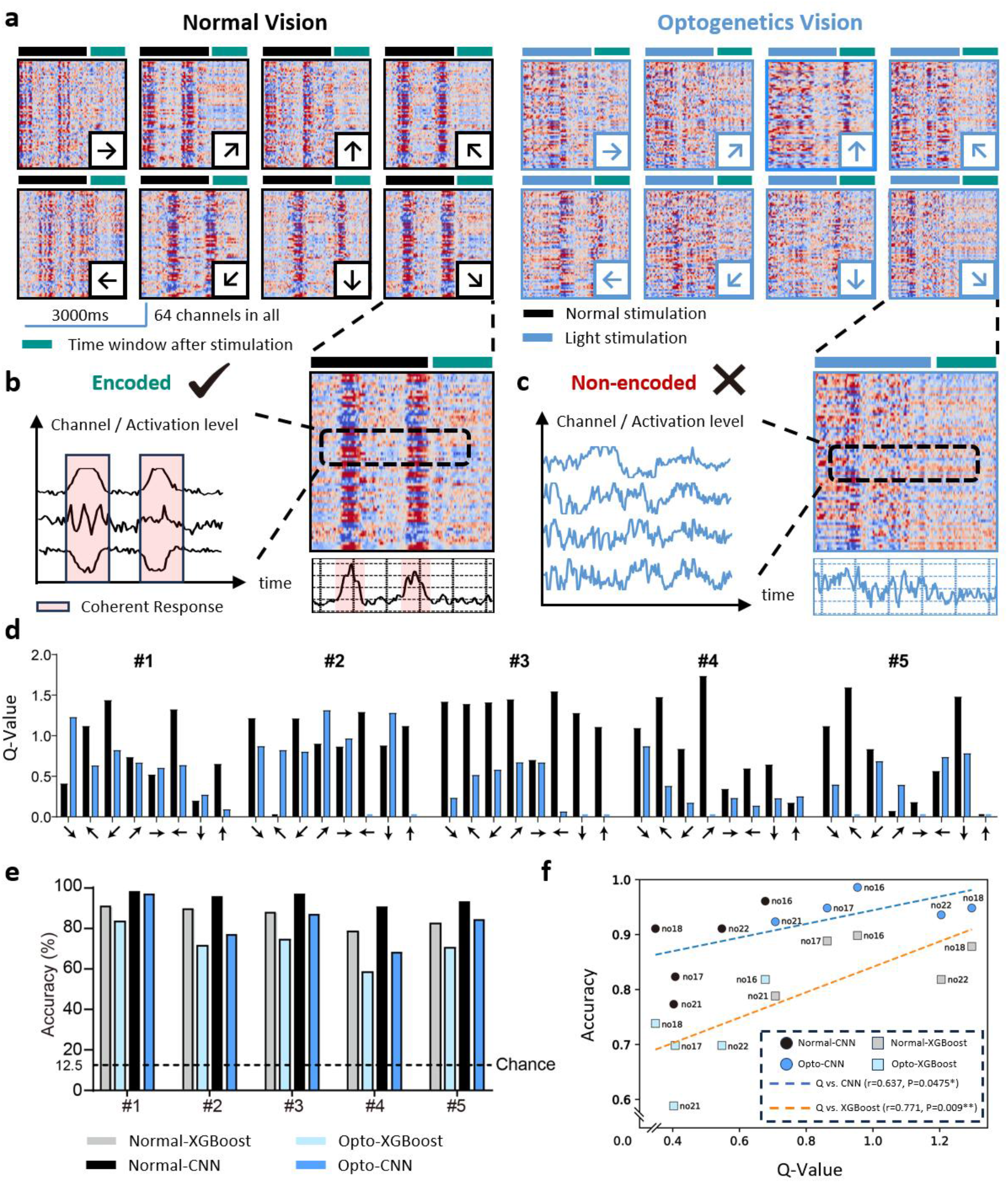
Artificial vision disrupts the temporal organization of direction-selective representations. **a,** Representative second-layer 1D-CNN feature maps from V1 responses to drifting gratings moving in eight directions under natural vision and optogenetic artificial vision. Each heat map contains 64 model-derived feature channels. Arrows indicate motion direction. Black and blue bars indicate the on/off periods of natural visual stimulation and optogenetic light stimulation, respectively; green bars mark the analysis window. Scale bar, 2,000 ms. Opto denotes optogenetic stimulation. **b,c,** Schematic and enlarged examples of encoded (**b**) and non-encoded (**c**) temporal feature patterns. The dashed boxes indicate the selected feature channels and time windows shown below. **d,** Double-peak quality score, Q, across motion directions for five mice. Higher Q values indicate clearer and more temporally organized double-peak feature patterns. Black, natural vision; blue, optogenetic stimulation. **e,** Direction-decoding accuracy using XGBoost and CNN classifiers under natural and optogenetic visual conditions. The dashed line indicates chance level for eight-direction classification. **f,** Relationship between mean Q value and direction-decoding accuracy. Points indicate animal-condition pairs, and dashed lines show linear fits for each classifier.

To quantify this difference, we defined a double-peak quality score, Q, which measured the salience, balance, and temporal regularity of the two-peak feature pattern. Across animals and directions, natural vision generally produced higher Q values, whereas optogenetic stimulation yielded lower Q values in most cases. This confirmed that the organized temporal feature pattern observed under natural vision was less consistently preserved under artificial vision (Fig. 5d).

We then tested whether this reduction in feature organization translated into weaker direction decoding. Using both an XGBoost classifier and a CNN decoder, motion direction was decoded more accurately under natural vision than under optogenetic stimulation across animals (Fig. 5e). This difference is consistent with the feature-level observations: under natural vision, direction-related features were more temporally organized and more separable, whereas under artificial vision, the weakened feature structure reduced discriminability. The relationship between feature organization and decoding performance was further supported by a positive correlation between mean Q values and classification accuracy across animals and visual states (XGBoost: r = 0.771, p = 0.009; CNN: r = 0.637, p = 0.047; Fig. 5f). This association suggests that stronger double-peak organization accompanies more reliable direction decoding.

Together, these results show that artificial vision retains motion-related signals in V1 but represents them in a less organized and less separable feature space than natural vision. Combined with the direction remapping observed in Section C, this indicates that the main limitation of dynamic artificial vision is not a simple loss of temporal tracking fidelity. Rather, artificial input generates a spatial-motion representation whose direction identity, gain structure, and learned feature organization differ from those produced by natural visual circuits. This provides a mechanistic basis for why downstream circuits may struggle to extract stable dynamic percepts from artificial visual input.

## Discussion

### After a century of pixel-printing: the cortex reorganizes artificial input rather than reproducing it

Nearly a century ago, neurosurgeons stimulating the human occipital lobe reported that patients perceived discrete spots of light ^32,33^. Subsequent work established an orderly retinotopic correspondence between cortical location and perceived visual-field position, such that focal cortical stimulation evokes phosphenes at predictable locations ^2,34,35^. This mapping licensed an intuitive engineering premise that has guided the field ever since. To restore vision, one would only need to "print" an image onto the cortex pixel by pixel, placing each stimulation site at its retinotopically matched position ^36,37^. Half a century of effort has shown that this premise fails. Rather than summing into a coherent image, neighbouring phosphenes interfere and coalesce, and the result is a blurred, unstable percept rather than the intended pattern ^6,38^. The field has gradually come to recognize that the cortex is not a display panel ^39^. This conclusion, however, was reached empirically, through the accumulation of failed attempts, and not from any direct measurement of what the cortex actually does with artificial input. Why the pixel-printing assumption fails has therefore remained unresolved.

Our within-subject, cross-stage framework was designed to measure this directly. In the same cortical substrate, we recorded responses first under intact natural vision and again under artificial vision. In the artificial condition, patterned light drives retinal ganglion cells directly and thereby bypasses the photoreceptor and inner-retinal circuitry that natural vision traverses (Fig. 1). Because stimulus content was matched across the two states, this design isolates the single variable that pixel-printing schemes implicitly ignore, namely the presence or absence of natural retinal encoding, and it lets us observe rather than infer how cortical representation responds. Two concrete observations followed. In the temporal domain, the geometry of V1 population states reorganized. Rates that natural vision compressed into overlapping, indistinguishable clusters became separable under artificial drive (Fig. 2). In the spatial domain, direction selectivity was preserved but its assignment was overturned, with preferred directions shifting systematically toward orthogonal axes rather than remaining fixed (Fig. 4). In neither dimension did the cortex simply reproduce the input at its retinotopically matched coordinates. Instead, it imposed its own reorganization. This locates the failure of pixel-printing more precisely than spatial cross-talk between electrodes can, because the cortex never operated as a point-to-point location map to begin with. It reads a spatiotemporal encoding structure that natural vision supplies and that the artificial pathway strips away. When identical stimulus content is delivered without that structure, it is reorganized rather than faithfully rendered. Whether this reorganization is uniform across dimensions, and what it implies for dynamic vision, requires a closer examination.

### A dissociable spatiotemporal gap: defining the bottleneck of today’s dynamic artificial vision

That reorganization, it turns out, is not uniform. Over the past five years, the field has moved beyond static pixel-printing. Using dynamically steered, temporally patterned stimulation, landmark studies ^3,4^ reliably elicited recognizable forms such as letters in both sighted and blind subjects ^40,41^. These studies established a pivotal idea that we build on directly: restoration hinges on dynamic codes that lead the cortex to actively interpret artificial input, rather than on brute-force spatial reproduction ^9,42^. These are genuine advances. Their dynamics, however, are trajectories traced over a stable phosphene topography, which amounts to a learned spatial coordinate transformation ^43–45^. This is categorically different from the dynamics of natural scenes, in which the content itself evolves in space and time ^9,46^. Whether artificially induced dynamic percepts and naturally evoked dynamic perception share a common cortical representation had not been testable, because no framework could compare them within one cortical substrate under matched input. This is the gap our design addresses ^41,47,48^.

Applying matched dynamic stimuli within this framework revealed an answer that is not uniform but dissociable across dimensions. In the temporal dimension, artificial vision did not merely preserve fidelity but extended it. Because the pathway bypasses the retinal low-pass stage ^49^, V1 sustained stimulus-locked entrainment across a broader frequency band than natural vision ^50^, which lost its temporal tuning at higher rates (Fig. 3). The relieved population-state compression seen in Fig. 2 therefore reflects a genuine gain in temporal tracking rather than an artefact. In the spatial dimension, the result was the reverse. The orthogonally remapped direction representations did not simply relocate information into a rotated but recoverable frame. Instead, the temporally organized feature structure that carries direction identity was itself eroded. Its double-peak organization weakened, and decodability fell accordingly, with feature quality tracking classification accuracy (XGBoost r = 0.771; CNN r = 0.637) (Fig. 5). This dissociation is the central finding of our study. Even with temporal fidelity fully preserved, and indeed enhanced, the spatial content of artificial input collapses into a poorly decodable representation. This directly answers a question that the recent dynamic-encoding paradigm could pose but not resolve, namely why escalating the strength, density, and temporal precision of stimulation has not yielded more naturalistic dynamic perception ^6,38^. The limiting factor is not how much dynamic signal is delivered, but whether its spatial structure survives in a form the cortex can read.

### An integrated model: from present mechanism to the future of prosthetic design

Having established that bypassing retinal encoding preserves temporal fidelity while collapsing spatial structure, we now ask what this dissociation means for the design of future visual prostheses. Fig. 6 integrates the results of Figs. 2–5 into a single conceptual model, using a naturalistic dynamic scene, a burst of fireworks (Fig. 6a, b), as a worked example. Such a scene is at once a distribution of local temporal frequencies and a field of local motion directions ^42^, so it loads both encoding axes at once. Rendering it through the natural and the artificial pathway therefore shows, in one picture, where an artificial code succeeds and where it fails, and why.

**Figure 6.**
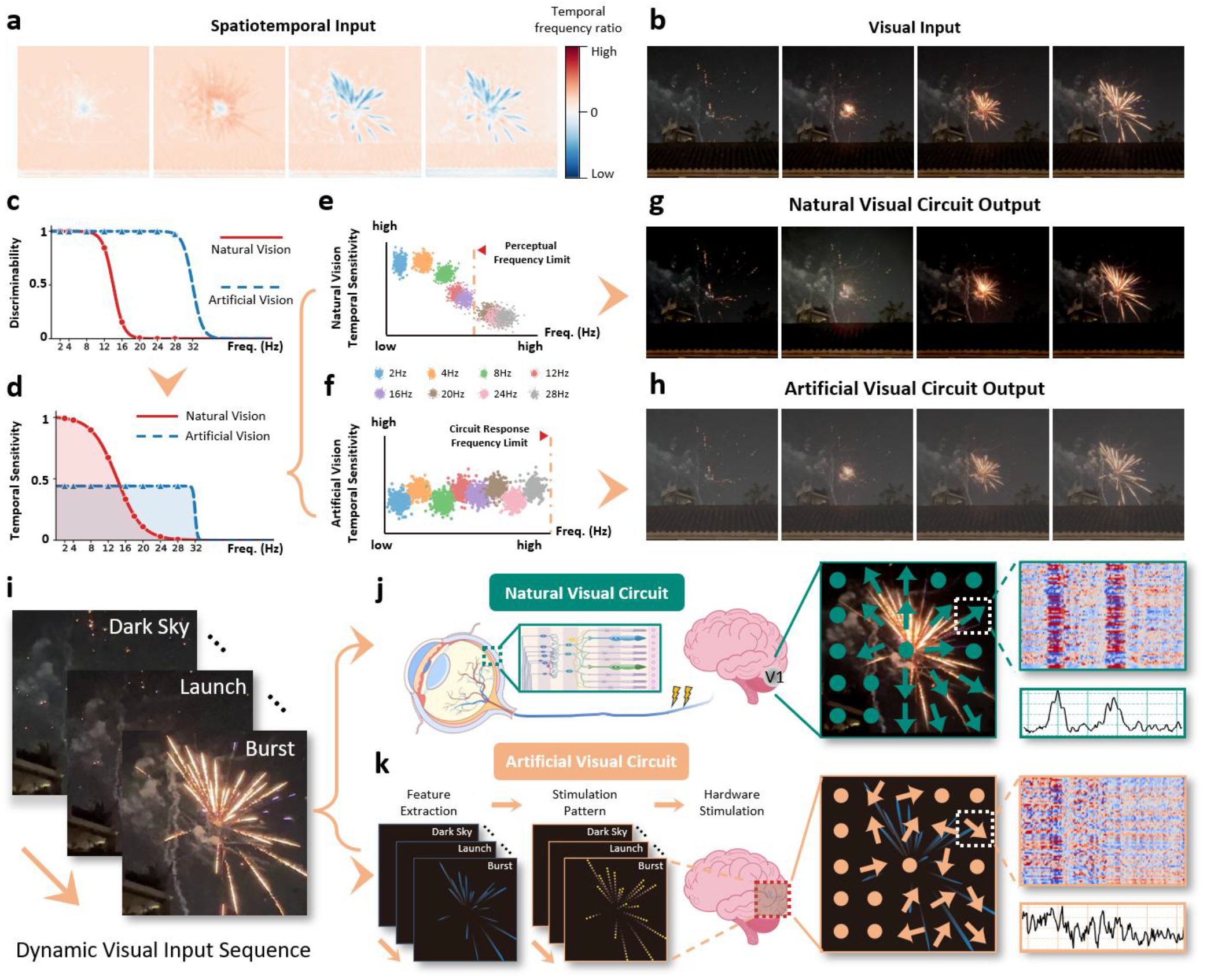
Integrated model of spatiotemporal dissociation and its implication for prosthetic encoding. **a**, Spatiotemporal input structure of a dynamic visual sequence (firework bursts); color indicates local temporal frequency ratio. **b**, Representative frames of the natural visual input. **c**, Frequency-discriminability curves derived from population-state separability (t-SNE, Sections A–B). **d**, Temporal sensitivity across frequencies, converted from **c**, reflecting encoding-resource allocation under each visual state. **e–f**, The two visual states resolved separately from **d**. **e**, Natural vision, where high frequencies receive reduced sensitivity below the perceptual threshold, defining a perceptual frequency limit. **f**, Artificial vision, where sensitivity is uniformly distributed and the ceiling is set by the circuit response frequency limit. **g–h**, Visual outputs arising from the two frequency-allocation strategies. **g**, Natural encoding concentrates resources on lower-frequency, behaviorally relevant components, forming a frequency-based attentional center; the firework coincides with this center and is enhanced while other components are attenuated. **h**, Artificial encoding, lacking upstream retinal processing, is frequency-decentralized, assigning uniform weight across components. **i**, Frames of a dynamic firework sequence spanning distinct temporal phases (dark sky, launch, burst). **j–k**, Schematic comparison of the two visual pathways. **j**, The natural visual pathway, from retinal encoding through upstream processing to V1, yielding accurate direction tuning and regular feature responses. **k**, The artificial visual pathway, in which features are extracted from the visual scene, converted into stimulation patterns, and delivered to the brain through the stimulating device; lacking upstream biological processing, it produces disrupted direction selectivity. Schematics contrast the two pathways and illustrate the proposed basis for the spatiotemporal dissociation observed in this study.

Along the temporal axis, we recast our discriminability measurements (Fig. 2) as a single frequency-discriminability profile contrasting the two pathways (Fig. 6c). Because attentional resources are finite ^51^, this profile implies a conserved allocation of sensitivity (Fig. 6d): discriminability at one band is withdrawn from another. Separating the two conditions yields two neural-representation models (Fig. 6e, f) whose ceilings have different origins, perceptual for natural vision and set by the circuit response limit for artificial vision. Processing the same input through each produces opposite allocations. Natural vision spends its budget on the low, behaviourally relevant frequencies and lets high-frequency detail fall away, forming an attentional centre on which the salient event is amplified and the background suppressed (Fig. 6g) ^52,53^. Artificial vision, lacking this upstream weighting, distributes sensitivity almost uniformly and treats every frequency as equally important (Fig. 6h). Whether the cortex can read such a decentralized signal is not settled, but our data give grounds for cautious optimism: V1 responds to high-frequency artificial drive with a regular, structured pattern (Fig. 3), suggesting it retains some capacity to represent this input and could adapt to it through plasticity. This adaptive potential, however, is not a licence to deliver input flat. The design lever is clear. A prosthetic front-end should not maximize temporal bandwidth, but reinstate the retina’s compressive, salience-forming filter ^54^, so that input arrives already organized around an attentional centre and thereby matched to the way the cortex decodes it.

Along the spatial axis, the collapse we quantified (Fig. 5) traces back to what each pathway computes before V1 (Fig. 6j, k). Under natural vision, moving structure passes through the full retinal cascade, where direction is computed and assigned before the signal reaches the cortex ^55,56^, and V1 inherits the accurate, organized direction tuning of natural stimulation (Fig. 6j). Under artificial drive, features are extracted from the scene and converted directly into stimulation patterns, so the retinal computations that establish direction are skipped entirely; V1 receives input that never underwent this encoding, which is why its direction tuning is remapped rather than reproduced (Fig. 4) and why the double-peak feature structure that carries direction has largely collapsed (Fig. 6k). Our directional stimulation is analogous to the rolling stimulation used to render letters, and for such sparse targets a fixed retinotopic correspondence is enough for the cortex to interpret the pattern without any rich internal directional structure. Even then, however, the underlying spatial representation remains collapsed, and this reliance on retinotopic correspondence rather than representational accuracy may be one reason such dynamic-stimulation approaches lose traction as they are extended toward complex, naturalistic scenes. As a scene approaches that complexity, and its meaning comes to reside in coherent motion, the missing structure becomes the binding constraint: the cortex is asked to synthesize motion from a drive that no longer carries an organized directional scaffold. This also marks the boundary of brain–machine mutual learning ^57,58^. The cortex can adapt to a novel code so long as the code is well structured, but once the scaffold itself has collapsed there is little stable structure left to learn from ^59^.

Fig. 6 thus makes a point that no single result could make alone: both axes fail from the same absent retinal stage, which in time relieves a compression that carried salience and in space dismantles a scaffold that carried direction. We cannot yet solve this directly. But the framework established here offers a route toward it, as a testbed in which stimulation algorithms can be trained and then validated at the level of the neural representation they evoke, rather than by perception alone ^60^. The goal is correspondingly specific: rather than delivering more signal, the encoding the retina would otherwise perform should be reconstructed at the point where stimulation is generated, so that the input is already compatible with cortical representation before it arrives.

## Methods

### Experimental Animals

A total of five male C57BL/6J mice, aged 2-4 months at the onset of the experiments, were used in this study. The animals were obtained from the Westlake University Animal Center. All mice were housed in a specific-pathogen-free (SPF) facility under highly controlled environmental conditions, maintaining a temperature of 22±1 ℃ and a relative humidity of 50±10%. The animals were kept on a standard 12-hour light/12-hour dark cycle to maintain natural circadian rhythms, with ad libitum access to standard rodent laboratory chow and sterilized drinking water. Prior to any experimental interventions, the animals were allowed to acclimatize to the housing environment for at least one week to minimize handling-induced stress. All experimental procedures, surgical protocols, and postoperative care were strictly conducted in accordance with the ethical guidelines for animal research and were formally reviewed and approved by the Westlake University Institutional Animal Care and Use Committee (IACUC, approval #: 24-017-MS-6).

### Electrode Implantation and Head-Post Surgery

Mice were anesthetized via intraperitoneal injection of 1.25% Avertin (200 µL/10 g) and secured in a stereotaxic frame with body temperature maintained at 37°C. Following a small craniotomy, a 16-channel microwire electrode array was implanted into the left primary visual cortex at stereotaxic coordinates AP −3.3 mm, ML +2.3 mm, and DV −0.46 mm relative to the cortical surface, alongside a reference screw placed over the contralateral cerebellum. During the application of dental cement to secure the electrode assembly, a custom head-post was simultaneously embedded and affixed to the skull. This head-post was utilized to immobilize the animal’s head in a fixed posture for viewing visual screen patterns during subsequent electrophysiological recordings^61^. Postoperatively, mice received 0.5% Meloxicam for 7 consecutive days for analgesia and fully recovered before experimental use.

### Establishment of Retinal Degeneration Model

To chemically induce retinal degeneration, N-methyl-N-nitrosourea (MNU) was utilized. Because MNU is light-sensitive and unstable in aqueous solutions, the compound was freshly dissolved in sterile physiological saline in a light-protected environment to achieve a final concentration of 4 mg/mL immediately prior to use. Each mouse was precisely weighed to ensure adequate dosing, followed by a single intraperitoneal (i.p.) injection of the MNU solution at a dosage of 60 mg/kg body weight. After the injection, the animals were returned to their standard housing conditions. This established protocol selectively induces rapid apoptosis of the outer retinal photoreceptor cells, resulting in a reliable model of profound blindness typically within 7 days post-injection.

### Viral Vectors and Optogenetic Transduction

To confer light sensitivity to retinal neurons following MNU-induced degeneration, mice received bilateral intravitreal viral injections. Under deep anesthesia, topical proparacaine and compound tropicamide were applied for ocular anesthesia and pupillary dilation. Using a 5 µL Hamilton microsyringe under a stereomicroscope, 2 µL of AAV2-Syn-hChR2(H134R)-mCherry (1.2 × 1013 vg/mL) was injected into each eye. This specific construct utilizes the Syn promoter to target the expression of channelrhodopsin-2 (ChR2) and the mCherry reporter primarily to retinal ganglion cells (RGCs). Injections were delivered slowly over 1– 2 minutes, and the needle was kept in place for an additional minute to prevent viral reflux and intraocular pressure fluctuations. Postoperatively, antibiotic eye ointment was applied. All subsequent experiments were conducted 28 days post-injection to ensure stable optogenetic protein expression.

### Retinal Tissue Preparation and Immunohistochemistry

To evaluate retinal morphology and viral expression, mice were euthanized and the eyes were enucleated. The retinas were carefully isolated, fixed in 4% paraformaldehyde (PFA), and cryoprotected in a sucrose gradient. The tissues were then embedded in optimal cutting temperature (OCT) compound and sectioned vertically into 20 µm-thick slices using a cryostat. For immunohistochemical staining, the retinal sections were first blocked and then incubated with a primary antibody against Glial Fibrillary Acidic Protein (GFAP) to selectively label Müller cells and astrocytes, followed by incubation with an Alexa Fluor 488-conjugated secondary antibody. The adeno-associated virus-transduced retinal ganglion cell were directly identified utilizing their inherent mCherry red fluorescence. Finally, the sections were counterstained with DAPI to visualize cell nuclei, mounted with an anti-fade mounting medium, and imaged using a fluorescence or confocal microscope.

### Statistical Analysis

Data distributions are visualized using Tukey box plots, where the central line indicates the median, the edges of the box represent the interquartile range (IQR, 25th to 75th percentiles), and the whiskers extend to the most extreme data points within 1.5 × IQR. Statistical analyses were performed to evaluate the differences between the normal vision group and the optogenetic vision (opto vision) group. Comparisons between these two groups were conducted using an independent two-tailed Student’s t-test. The threshold for statistical significance was defined as *p < 0.05. In all figures, significance levels are denoted using the following conventions: *p < 0.05, **p < 0.01, and ***p < 0.001.

### Electrophysiological recording and signal acquisition

Electrophysiological recordings were performed in awake head-fixed mice bearing chronically implanted electrodes. All implantation surgeries were completed before the recording experiments, and the implanted head connector was used to interface the electrodes with the recording system. Recordings were initiated no earlier than 7 days after surgery and only after signal stability had been confirmed by continuous monitoring of waveform shape and baseline noise across channels. Neural signals were routed through the head connector to a 16-channel amplifier (Plexon DigiAmp, Plexon), amplified at 2000× gain, band-pass filtered at 300–5000 Hz, and acquired using an OmniPlex Neural Recording Data Acquisition System (Plexon) at a sampling rate of 40 kHz^62^. External digital high/low logic signals were delivered to the acquisition system via a BNC interface and recorded at 1 kHz for offline synchronization with neural activity. Spike extraction was performed offline using Plexon Offline Sorter. Continuous signals were first high-pass filtered with a 250 Hz Bessel filter. Threshold-crossing spikes were then detected using the standard deviations from mean of peak heights histogram method, with the threshold set at 3.0 sigma. The extracted threshold-crossing spikes were used for subsequent analyses.

### Visual function assessment and receptive field mapping

Before formal experiments, mice with normal vision were screened for visually evoked neural responses and receptive field location. Awake head-fixed mice were positioned with the tested eye facing a display monitor (220 mm × 140 mm) at a distance of 140 mm, corresponding to approximately 76.3° × 53.1° of visual angle. The display was divided into a 4 × 4 grid of 16 rectangular sectors. Black-and-white drifting square-wave gratings were presented pseudo-randomly within each sector, with each sector stimulated 30 times. Gratings were presented at a direction of 0° for 2 s using multiple drift speeds. Visually evoked responses were assessed from firing rates during stimulus presentation. Mice showing reliable grating-evoked activity were included in subsequent experiments. Receptive field location was estimated from the spatial distribution of firing rates across the 16 sectors, and the monitor position was slightly adjusted so that the receptive field was centered on the display for subsequent visual stimulation.

### Visual stimulation setup for normal vision

Visual stimuli under normal vision were generated by a small color LCD monitor, using the same general configuration as described for visual function assessment. Awake head-fixed mice were positioned with the tested eye facing the monitor. The monitor had an active display area of 220 mm × 140 mm, operated at 60 Hz, and was typically placed 140 mm from the tested eye, corresponding to approximately 76.3° × 53.1° of visual angle. The monitor position was slightly adjusted for individual mice based on receptive field mapping to center the receptive field on the display.

### Optogenetic stimulation setup

Optogenetic stimulation was delivered using a 473 nm blue LED array. The LEDs were commercial blue LEDs (LXML-PB0I, LUMILEDS), each with a diameter of approximately 3.5 mm. The array consisted of 8 × 8 LEDs uniformly arranged and mounted within a 100 mm × 100 mm area on a printed circuit board. Each LED was independently controlled by an STM32F103C8T6 microcontroller through decoder and driver circuits. The size and position of the LED stimulation array were matched to the normal visual stimulation setup and adjusted according to the receptive field center of each mouse. The stimulus irradiance under this setup was controlled manually and measured with a photometer around approximately 5 mW/cm²^19,63^.

### Stimulus paradigms for temporal frequency and spatial tuning

To investigate the modulation of temporal coding, two visual stimulation paradigms were used in this study: drifting gratings with different temporal frequencies (Fig. 2) and SSVEP stimulation at different frequencies (Fig. 3). For the drifting grating experiment, black-and-white drifting square-wave gratings were presented on the display with a drift direction of 315°. The stimulus area was matched to the LED array size, both measuring 100 mm × 100 mm, and was positioned 140 mm from the tested eye, corresponding to a visual angle of approximately 39.3° × 39.3°. Each grating contained four complete black-white cycles across the stimulus width, resulting in a spatial frequency of approximately 0.10 cycles/degree. Gratings were presented at eight temporal frequencies: 2, 4, 8, 12, 16, 20, 24, and 32 Hz. Each frequency was presented for 50 trials in a pseudo-random order, with each trial lasting 2 s. After each stimulus presentation, mice were given a 2 s inter-trial interval, during which a gray background was displayed on the monitor and all LEDs in the LED array were turned off. Stimulus timing was synchronized with neural recordings using digital triggers^61^.

For the SSVEP stimulation experiment, stimuli were delivered by the LED array using a centrally symmetric cross-shaped pattern within the same 100 mm × 100 mm stimulation area. The pattern was presented with on/off luminance modulation at eight flicker frequencies: 0.5, 1, 2, 4, 8, 12, 16, and 32 Hz. Each frequency was repeated for 50 trials in a pseudo-random order. Each trial lasted 2 s and was followed by a 2 s inter-trial interval, during which all LEDs were turned off.

### t-SNE Clustering

For each mouse and each experimental condition, t-SNE was performed independently using trial-wise spike count feature vectors. For each trial, spike counts were calculated for each recorded channel in 200 ms bins from 200 ms before to 2.0 s after the stimulus trigger, and the counts from all channels and time bins were concatenated into one feature vector. t-SNE was applied to these feature vectors with two output dimensions. The resulting two-dimensional embeddings were plotted separately for the normal vision and optogenetic stimulation conditions, with points colored according to stimulus label. Within each label, Mahalanobis distance was calculated in the two-dimensional t-SNE space using robust covariance estimation, and the 50% most central trials were displayed. Because body movements during awake recordings caused large trial-to-trial fluctuations in neural firing, this filtering step was used only to visualize the main response trend and did not affect the quantitative analyses.

### XGBoost

XGBoost was used as a baseline decoding model to classify stimulus conditions from trial-wise neural population responses^64^. For each trial, spike counts were calculated for each recorded channel within the time window from 0.2 s before to 2.0 s after stimulus onset, using 0.2 s time bins. To reduce the influence of differences in overall firing levels across channels, spike counts were normalized at the channel level. The normalized spike counts from all channels and time bins were then concatenated to form a population response vector for each trial. The XGBoost classifier was trained to predict stimulus labels from these response vectors, and model performance was evaluated using stratified cross-validation. Classification accuracy and F1-score were used as performance metrics.

### CNN model & Feature Map

A one-dimensional convolutional neural network was trained to decode stimulus labels from trial-wise spike activity^65^. For each trial, spike times from all recorded channels were aligned to the stimulus period and converted into a binary spike tensor with the shape of channels × time. Each time bin was assigned a value of 1 if at least one spike occurred in that bin and 0 otherwise. In the CNN analysis, each trial was represented by 1000 time bins over the analyzed stimulus window.

The CNN consisted of two temporal convolutional blocks followed by fully connected layers. The first convolutional layer mapped the input channels to 32 feature channels using a kernel size of 5, followed by ReLU activation and max pooling. The second convolutional layer mapped 32 feature channels to 64 feature channels with the same kernel size, followed by ReLU activation and max pooling. The resulting features were flattened and passed through a fully connected layer with 128 units, ReLU activation, dropout, and a final linear layer for stimulus classification.

For each experiment, trials were split into training and validation sets using an 80%/20% class-balanced split with a fixed random seed of 42. The model was trained using cross-entropy loss and the Adam optimizer with a learning rate of 1 × 10^-3. Training was performed for 50 epochs with a batch size of 32. After each epoch, validation accuracy was calculated, and the model parameters with the highest validation accuracy were retained.

To examine the temporal features learned by the CNN, activation maps from the second convolutional layer were extracted using the validation trials. A forward hook was applied to the second convolutional layer, and the resulting Conv2 activations were collected for each validation trial. For each stimulus class, Conv2 activations were averaged across validation trials belonging to that class, producing a class-averaged feature map. Representative activation waveforms were further obtained by averaging the class-averaged Conv2 feature map across feature channels and then standardizing the resulting temporal waveform to zero mean and unit variance for visualization.

### Direction Tuning and Clustering

To assess direction selectivity, drifting gratings were presented in eight directions (0°, 45°, 90°, 135°, 180°, 225°, 270°, and 315°). Each direction was repeated for 50 trials. For each recorded channel and each direction, spike counts were calculated during the stimulus presentation period and averaged across the 50 trials. The resulting mean spike count as a function of stimulus direction was plotted as a direction tuning curve for each channel^16,17^.

To quantify the similarity of direction tuning properties across experimental conditions, direction tuning curves from all recorded channels were first normalized using z-score transformation within each channel:

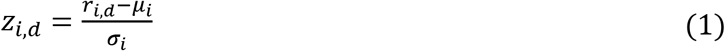

where *r_i_*_,*d*_ is the mean spike count of channel *i* in direction *d*, *μ_i_* is the mean spike count across all eight directions for channel *i*, and *σ_i_* is the standard deviation across directions.

Normalized tuning curves were then clustered into four groups using k-means clustering, performed independently for each experimental condition. Clusters were sorted by the number of channels assigned to each cluster and labeled A, B, C, D in descending order of cluster size. For visualization and correlation analysis, only clusters containing at least 2 channels were included. For each retained cluster, a cluster-averaged tuning curve was computed by averaging the normalized tuning curves of all channels assigned to that cluster.

To assess the stability of direction tuning across normal vision and optogenetic stimulation conditions, Pearson correlation coefficients were calculated between cluster-averaged tuning curves from the two conditions. The Pearson correlation coefficient between two tuning curves **x** and **y** was defined as:

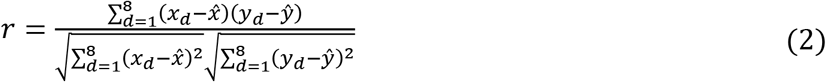

where *x_d_* and *y_d_* are the normalized spike counts at direction *d*, and *x̄* and *ȳ* are the means across all eight directions. For the same mouse, Pearson correlation coefficients were calculated between cluster-averaged tuning curves under normal vision and optogenetic stimulation conditions, resulting in a correlation matrix representing inter-cluster similarity (Figure 4c). Additionally, for each individual channel, the correlation coefficient was calculated between its tuning curve under normal vision and its tuning curve under optogenetic stimulation, yielding channel-level self-similarity measures (Figure 4d).

### Tuned and Non-Tuned Frequency Analysis

To quantify the strength of SSVEP encoding in CNN feature maps, we computed the percentage of power at the target SSVEP frequency relative to total power across all frequencies. For each of the 64 feature channels in the second convolutional layer, a 2 s time window was extracted from the class-averaged Conv2 feature map, corresponding to the duration of visual stimulus presentation. The extracted signal was denoted as *x*(*t*)where *t* ∈ [0,2]s. The power spectral density (PSD) was computed using Welch’s periodogram method:

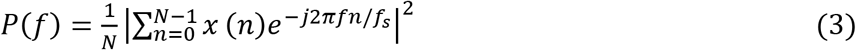

where *N* is the number of time points, *f_s_* is the sampling rate of the feature map (60 Hz for both normal and optogenetic conditions), and *f* is the frequency in Hz.

The target SSVEP frequency was defined as half of the stimulus frequency (*f*_SSVEP_ = *f*_stim_/2), reflecting the characteristic frequency-doubling response observed in V1 neurons. The SSVEP power ratio *R* was computed as the ratio of power within a ±0.5 Hz band centered at the target frequency to the total power across all frequencies:

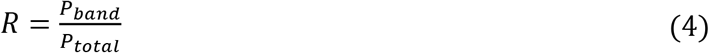

where:

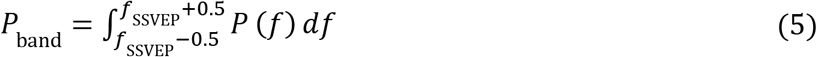

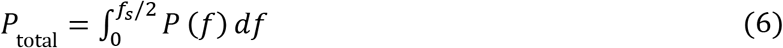

Feature channels were classified into two categories based on their power ratio values:

#### Tuned channels

*R* ≥ 0.3, indicating reliable SSVEP encoding

#### Not-tuned channels

*R* < 0.3, indicating weak or absent SSVEP encoding

For each experimental condition (normal vision and optogenetic stimulation) and each stimulus frequency, the distribution of power ratio values across all 64 feature channels was computed (Figure 3b). The number of tuned channels out of 64 total channels was quantified for each condition and frequency (Figure 3c).

### Double-Peak Quality Assessment in Representative Waveforms

To quantify the clarity of double-peak structures in neural response waveforms, we developed a double-peak quality metric. For each representative waveform of a direction class, a 2 s time window following stimulus onset was extracted, corresponding to the primary response period of neurons. The waveform was first smoothed using a Gaussian filter (σ = 2.0) to reduce high-frequency noise. Peaks were detected using the *scipy.signal.find_peaks* function with a minimum prominence of 0.8 and a minimum inter-peak distance of 200 ms to exclude spurious peaks caused by noise.

Among all detected peaks, the two peaks with the highest prominence values were selected as the primary peak pair P1 and P2, with P1 occurring earlier in time. Based on observations across all experimental data, samples exhibiting clear double-peak structures consistently showed an inter-peak interval of approximately 700 ms, reflecting the characteristic temporal dynamics of V1 neurons in response to motion stimuli. Therefore, the double-peak quality score was designed with an expected inter-peak interval of 700 ms and a tolerance range of ± 400 ms.

The double-peak quality score Q was computed as:

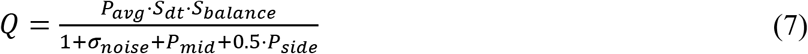

where:

*P*_avg_ = (*P*_1_ + *P*_2_)/2 is the average prominence of the two primary peaks, reflecting their salience

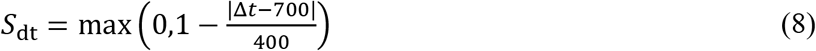

is the inter-peak interval score, where Δ*t* is the time interval (ms) between P1 and P2. The score approaches 1 when the interval is close to the observed typical value of 700 ms and decreases to 0 when the deviation exceeds 400 ms

*S*_balance_ = min(*A*_1_, *A*_2_)/max(*A*_1_, *A*_2_) is the height balance score, where *A*_1_ and *A*_2_ are the amplitudes of P1 and P2. This term rewards symmetric double-peak structures with similar peak heights
*σ*_noise_ is the standard deviation of the waveform, serving as an estimate of overall noise level
*P*_mid_ is the intermediate peak penalty. If a peak with prominence exceeding 0.6 exists between P1 and P2, then *P*_mid_ = max(0, *P*_intermediate_ − 0.6), where *P*_intermediate_ is the maximum prominence of intermediate peaks. This term penalizes unclear multi-peak structures
*P*_side_ is the outlier peak penalty. If peaks with prominence exceeding 1.0 exist before P1 or after P2, the penalty accumulates as *P*_side_ = ∑max(0, *P*_outlier_ − 1.0). This term is weighted by 0.5 in the denominator to reduce sensitivity to distant interfering peaks

Higher Q values indicate clearer double-peak structures with inter-peak intervals closer to the typical value of 700 ms and more symmetric peak heights. This metric comprehensively considers peak salience, temporal interval accuracy, structural symmetry, and the influence of interfering peaks and noise, enabling quantitative distinction between high-quality double-peak responses and noisy or irregular response patterns.

To investigate the relationship between double-peak response quality and direction discrimination performance, we computed the mean double-peak quality *Q̄* for each experimental session (5 mice × 2 conditions = 10 sessions):

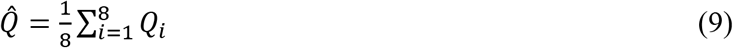

where *Q_i_* is the double-peak quality score for direction class *i*. A linear regression model was used to evaluate the relationship between mean double-peak quality *Q̄* and overall validation accuracy:

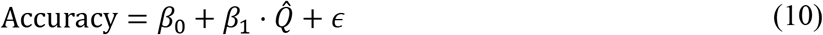

where *β*_0_ is the intercept, *β*_1_ is the regression slope, and *ε* is the residual term. Model goodness-of-fit was assessed using the coefficient of determination *R*^2^, and the significance of the correlation was tested using the Pearson correlation coefficient *r* and its associated p-value. Additionally, paired t-tests were used to compare mean double-peak quality and overall accuracy between normal vision and optogenetic stimulation conditions, assessing the impact of optogenetic disruption on double-peak response structure and direction discrimination performance. Statistical significance was defined as p < 0.05.

### Preferred direction and Mean response ratio

To assess the impact of optogenetic stimulation on direction selectivity, we computed the preferred direction shift (PD shift) and mean response ratio for each recording channel under normal vision and optogenetic stimulation conditions.

For each motion direction, 50 repeated trials were presented. For each channel, spike counts were computed within the stimulus presentation window [0, 2] seconds and averaged across the 50 trials to obtain the mean response for that direction. By computing the mean responses across 8 motion directions (0°, 45°, 90°, 135°, 180°, 225°, 270°, 315°), the channel’s direction tuning curve was constructed.

Preferred direction (PD) was defined as the direction with the maximum response in the tuning curve: where *R̄* (*θ*)is the mean spike count across 50 trials within the stimulus window [0, 2] seconds for direction *θ*.

PD shift quantifies the change in preferred direction for the same channel between normal vision and optogenetic stimulation conditions:

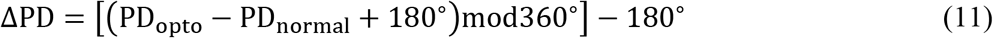

This formula ensures that the shift is constrained to the range [-180°, 180°], where positive values indicate clockwise shifts and negative values indicate counterclockwise shifts. PD shift reflects the impact of optogenetic disruption on neuronal direction selectivity: larger PD shifts indicate substantial alterations in the channel’s direction tuning properties.

Mean response ratio quantifies the impact of optogenetic stimulation on the overall response magnitude of neurons. For each channel, within the stimulus window [0, 2] seconds and averaged across 50 trials, the mean response across all 8 motion directions was computed separately for normal vision and optogenetic stimulation conditions:

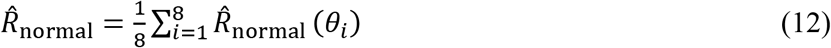

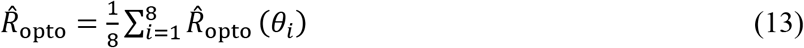

where *R̄*_normal_(*θ_i_*)and *R̄*_opto_(*θ_i_*)are the mean spike counts across 50 trials within the [0, 2] second window for direction *θ_i_* under normal vision and optogenetic stimulation, respectively.

The mean response ratio was defined as:

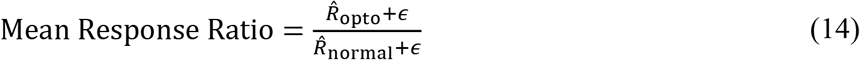

where *ε* = 10^−8^is a small constant to prevent division by zero. This ratio reflects the modulation of overall neuronal activity by optogenetic stimulation: ratio > 1 indicates enhanced responses, ratio < 1 indicates suppressed responses, and ratio ≈ 1 indicates minimal impact on overall response magnitude.

## Data availability

Data availability All data collected and analyzed in this study are available from the corresponding author upon reasonable request.

## Code availability

The code used to perform all analyses in this study is available from the corresponding author upon reasonable request.

## Funding

Brain Science and Brain-like IntelligenceTechnology-National Science and Technology Major Project (Grant No.2022ZD0208805).

The "Pioneer" and "Leading Goose" R&D Program of Zhejiang (2024C03002).

## Author Contributions

X.G., Y.Z. and C.W. conceived the concept. X.G. and W.R. designed the experiments. X.G., Y.Z., Y.X. and B.S. designed and built the experimental platform. X.G., W.R., B.Z., Y.Z., Y.X. and G.H. performed the experiments. X.G. carried out the data analysis. X.G. and C.C. designed the deep learning model. X.G. and W.R. wrote the manuscript. Y.C., J.Y. and M.S. provided guidance and supervised the project. All authors reviewed and approved the final manuscript.

---

**Figure S1.**
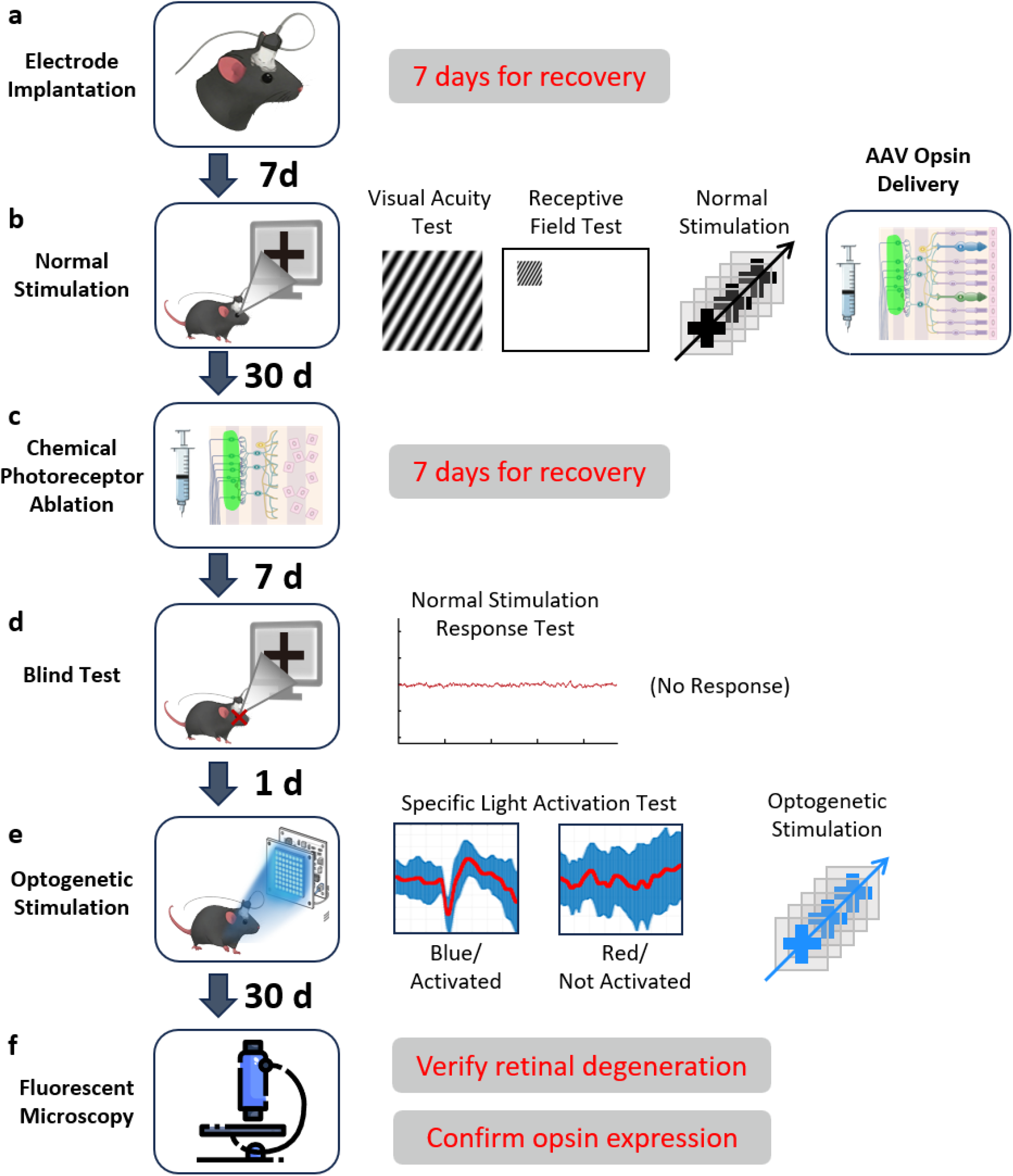
Experimental pipeline of the study. **a**, Microelectrodes are first implanted for stable neural recording, followed by a 7-day recovery period. **b**, Baseline visual functions, including visual acuity and receptive fields, are evaluated using display-based normal stimulation. Subsequently, AAV carrying opsin genes is delivered, and the animals are maintained for 30 days to allow for optimal viral expression. **c**, A specific chemical agent is injected to ablate the original photoreceptors, inducing retinal degeneration over a 7-day period. **d,** Functional blindness is then confirmed by the absence of electrophysiological responses to normal visual stimuli. **e**, During the optogenetic stimulation phase, targeted light projection (e.g., blue light) successfully elicits opsin-mediated neural responses, whereas control wavelengths (e.g., red light) fail to induce activation. **f**, After 30 days of functional testing, fluorescent microscopy is performed to verify the chemically induced retinal degeneration and confirm the successful expression of opsins.

**Figure S2.**
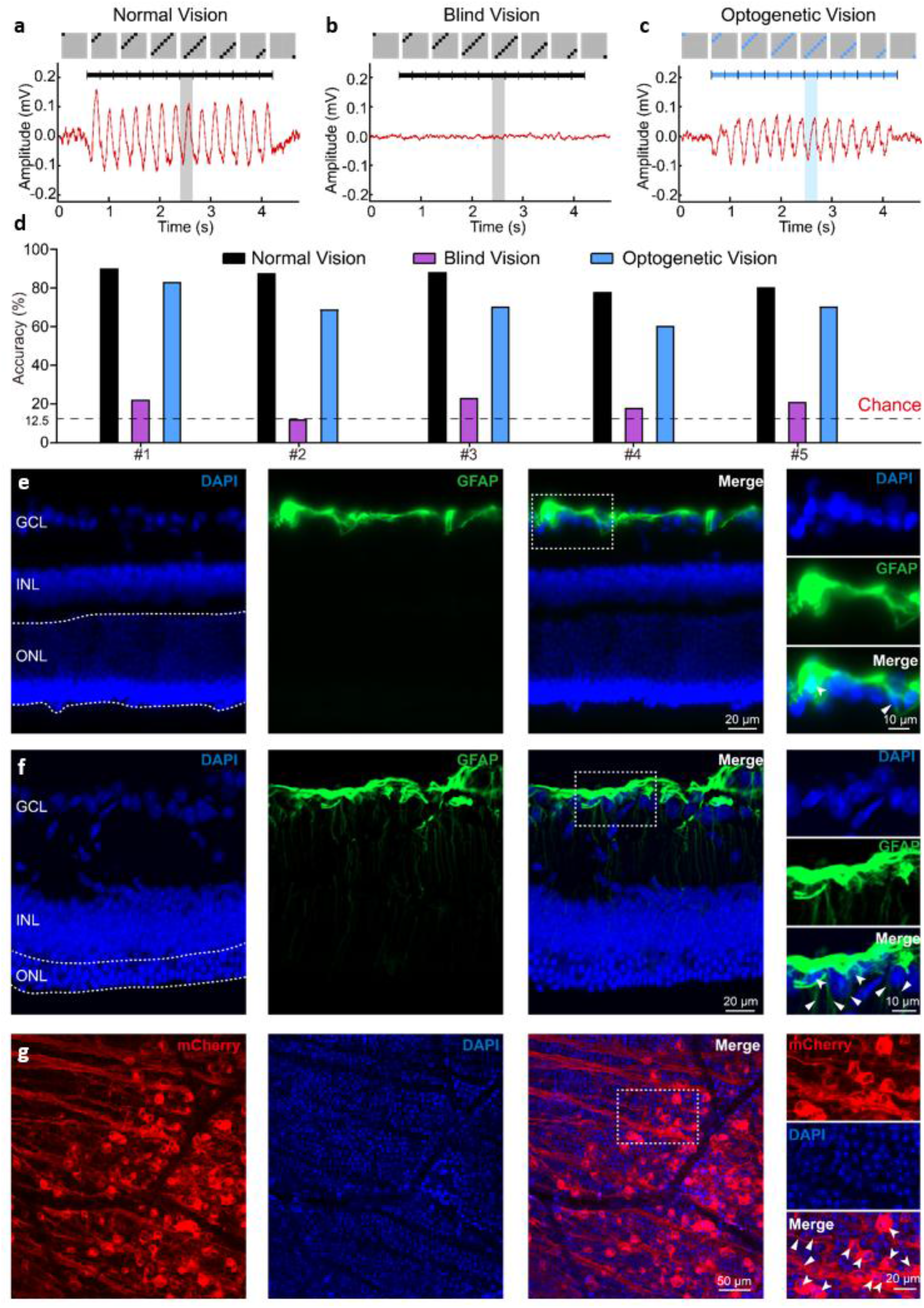
Validation of V1 neural responses and stimulus decoding across varying visual stages. **a-c,** Representative waveforms recorded from the primary visual cortex (V1) in response to a drifting grating stimulus across three longitudinal stages in the same subject: **a,** Normal vision, with the grating presented on an LCD monitor; **b,** Blind vision, with the same monitor stimulation post-blinding; and **c,** Optogenetic vision, with spatiotemporally equivalent LED pattern stimulation following successful opsin expression. Top panels illustrate the schematics of the visual stimuli. Horizontal lines denote the stimulus duration. Gray and blue shaded vertical blocks highlight a single oscillatory cycle. Note the robust stimulus-locked cortical responses in both normal and optogenetic conditions, which are completely absent in the blind condition. **d,** Decoding accuracy for grating directions across five individual mice (#1–#5) under the three visual conditions. The accuracy was evaluated using an XGBoost machine learning classifier based on the recorded V1 signals. Black, purple, and blue bars represent normal, blind, and optogenetic vision stages, respectively. The dashed line indicates the chance level (12.5%). **e-f,** Immunofluorescence confocal images of retinal cross-sections from **e**, a healthy control mouse and **f,** a blinded mouse. DAPI (blue) labels cell nuclei, indicating the retinal layers: ganglion cell layer (GCL), inner nuclear layer (INL), and outer nuclear layer (ONL). GFAP (green) labels glial cells. In the healthy retina **e**, GFAP expression is largely restricted to the GCL. In contrast, the blinded retina **f** exhibits structural disruption and severe reactive gliosis, evidenced by massive GFAP upregulation and glial processes extending radially through the retinal layers. Magnified inset panels correspond to the dashed white boxes. **g**, Confocal images of the retina demonstrating successful viral transduction. The mCherry-tagged opsin (red) is robustly expressed in retinal ganglion cells (RGCs). DAPI (blue) labels the cell nuclei. Magnified insets on the right highlight individual opsin-expressing RGC somas (white arrowheads).

**Figure S3.**
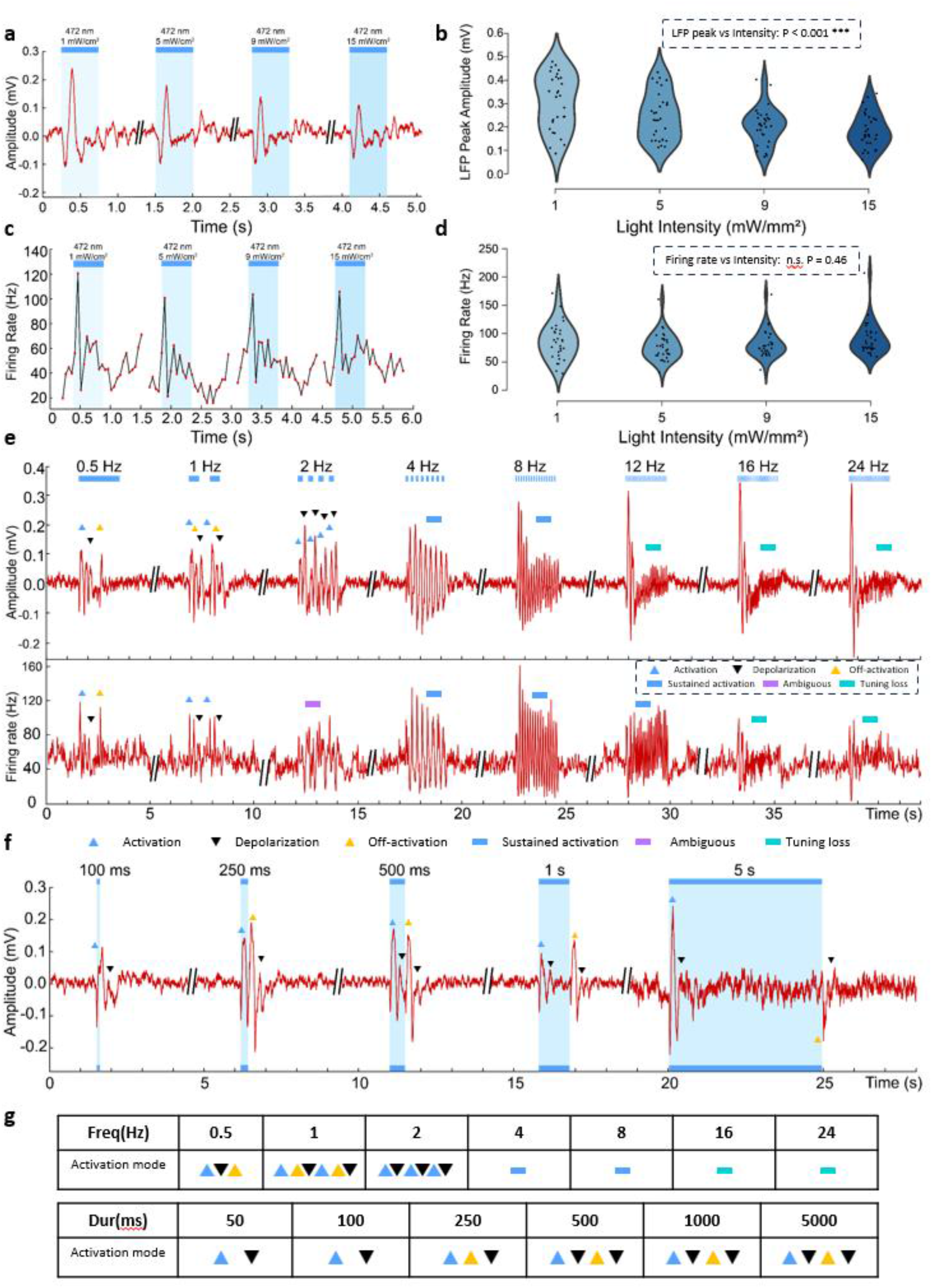
Characterization of V1 neural responses to varying optogenetic stimulation parameters. **a** Representative local field potential (LFP) traces recorded in V1 elicited by 472 nm blue light pulses at increasing intensities (1 to 15 mW/mm^2^). Light blue shaded regions indicate stimulus duration. **b**, Violin plots quantifying the peak LFP amplitude as a function of light intensity. LFP peak amplitudes significantly varied across intensities (P < 0.001). **c-d** Corresponding continuous firing rate profiles **c** and quantitative comparison of peak firing rates **d**, evoked by identical graded light intensities. No significant intensity-dependent modulation was observed for the firing rates (n.s., P = 0.46). **e,** Representative LFP (top) and firing rate (bottom) responses to optogenetic stimulation delivered at different temporal frequencies (0.5 to 24 Hz). Colored markers and bars denote distinct temporal response phases, including activation (blue upward triangle), depolarization (black downward triangle), off-activation (yellow upward triangle), sustained activation (light blue bar), ambiguous responses (purple bar), and tuning loss (teal bar). **f,** Representative LFP traces evoked by single continuous light pulses of varying durations (100 ms to 5000 ms), annotated with corresponding temporal activation phases. **g,** A summary table detailing the distribution and transitions of distinct activation modes across varying stimulation frequencies (top) and durations (bottom). Symbols correspond to the legend defined in **e** and **f**.

**Figure S4.**
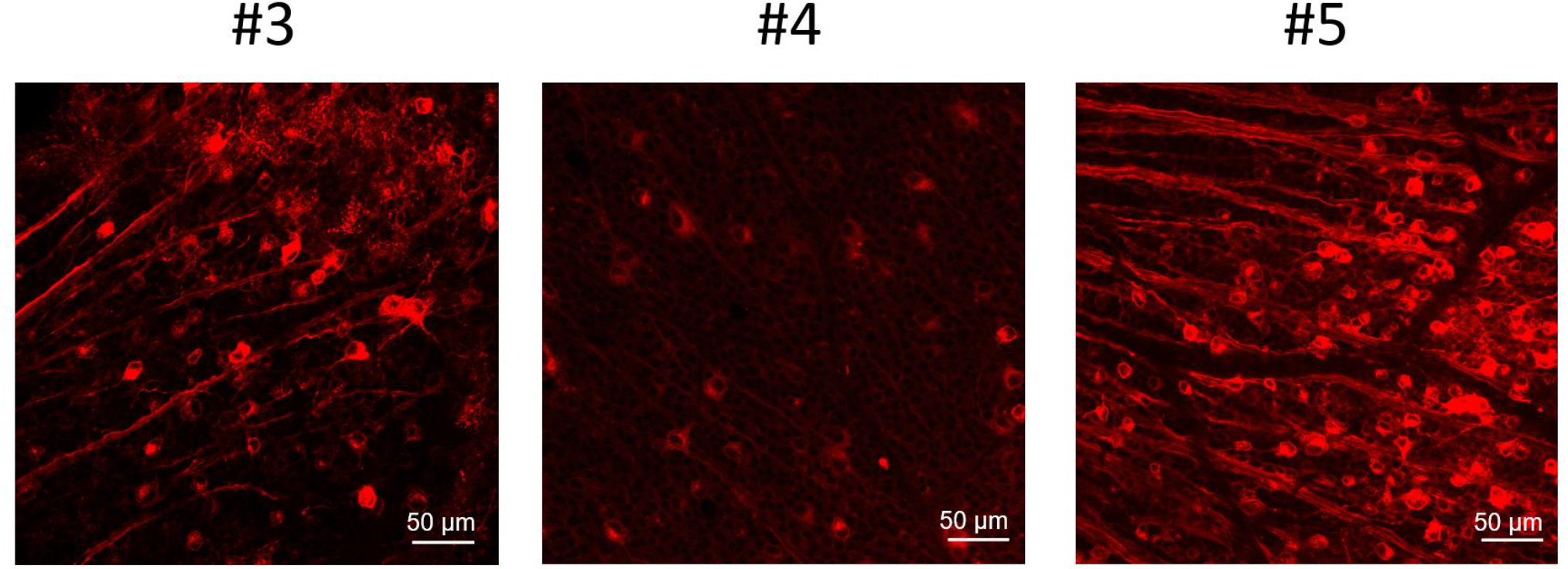
Inter-individual variability in opsin expression across subjects. Representative confocal images of mCherry-tagged opsin expression (red) in retinal ganglion cells from three mice (#3, #4, and #5) following identical viral injection doses and a uniform 30-day transduction period. The images illustrate intermediate (#3), low (#4), and high (#5) expression levels, demonstrating substantial inter-individual variability in transduction efficiency despite standardized procedures. Scale bars, 50 μm. Note: For illustrative purposes, the confocal image of mouse #5, which represents high expression levels here, is identical to the representative example provided in Figure S2.

**Figure S5.**
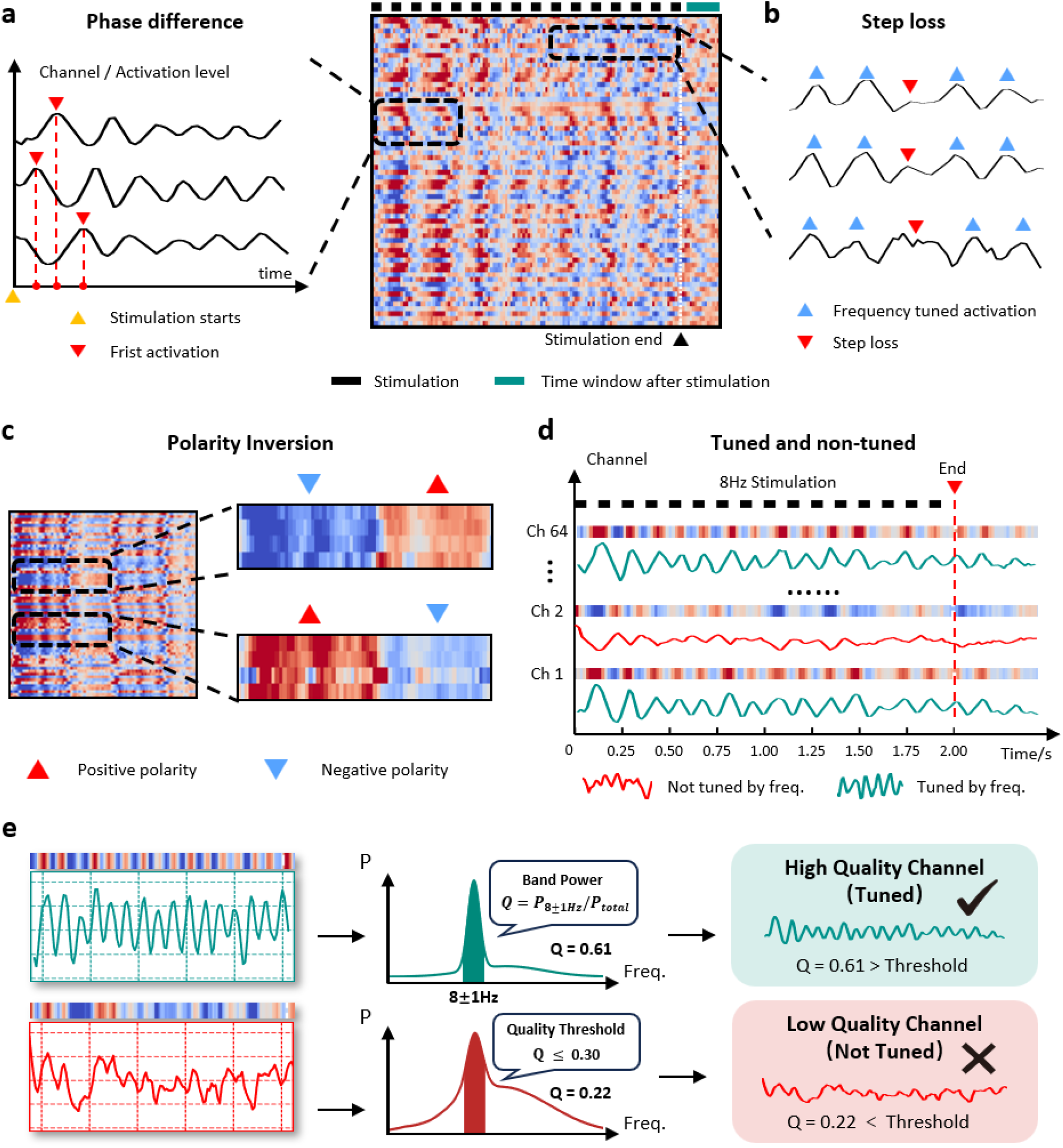
Characterization of CNN feature maps under SSVEP stimulation. **a**, Phase difference: temporal shift of the first activation peak (red triangles) relative to stimulus onset (yellow triangle) across channels. **b**, Step loss: missed activation cycles (red triangles) amidst entrained responses (blue triangles) at higher frequencies. **c**, Polarity inversion: switching between positive (red) and negative (blue) values reflects transitions between distinct representational states in the 1D-CNN feature space. **d**, Representative tuned (green) and non-tuned (red) channels under 8 Hz stimulation. **e**, Quantitative criterion for tuned-channel classification. The band-power ratio Q is defined as the spectral power within ±1 Hz of the stimulation frequency divided by total power. Channels with Q > 0.30 are classified as tuned (top, Q = 0.61); otherwise non-tuned (bottom, Q = 0.22).

**Figure S6.**
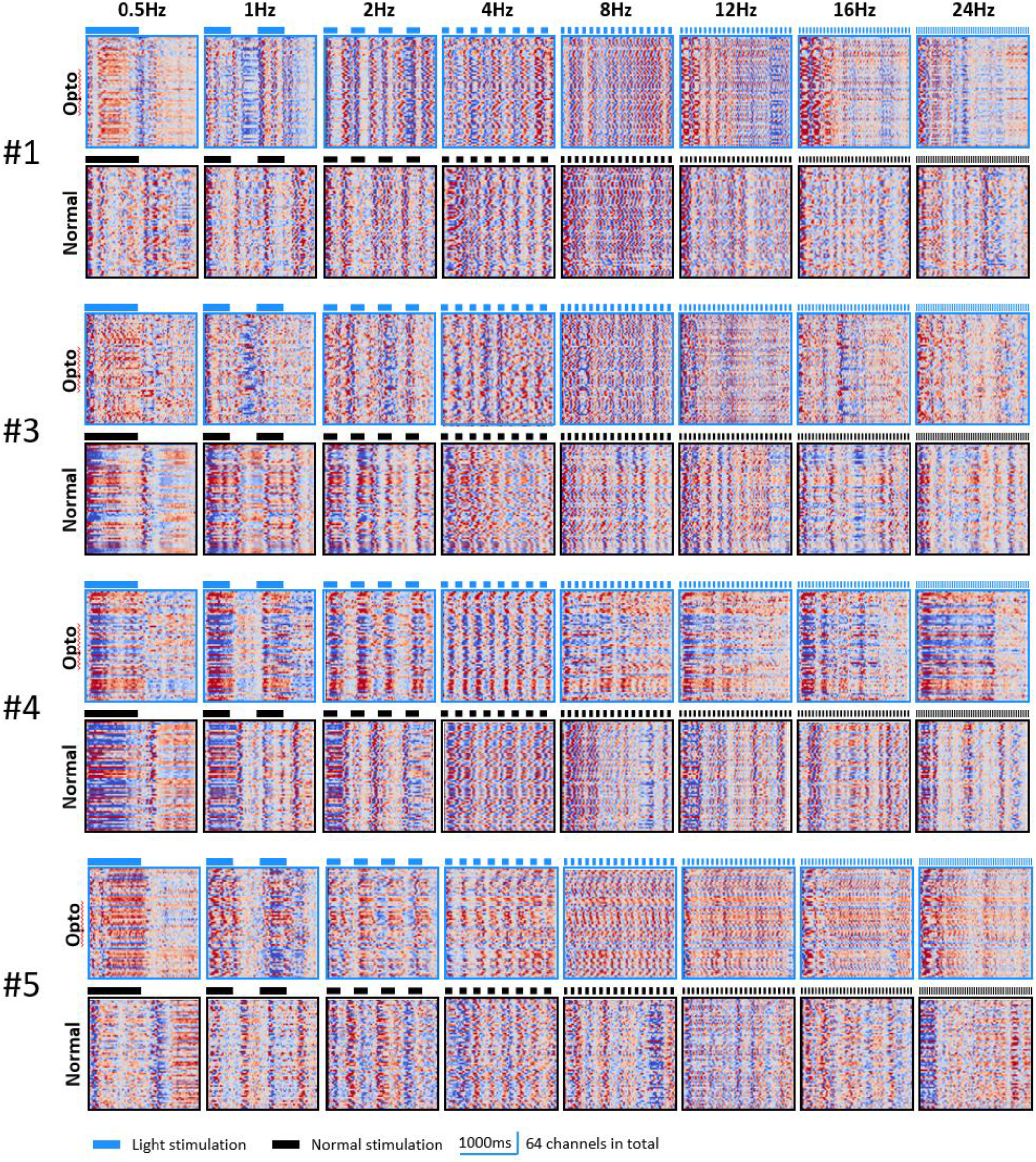
CNN feature maps of V1 responses across all subjects under optogenetic artificial vision and natural vision. CNN feature maps extracted from V1 responses to SSVEP flicker stimulation in individual mice (excluding subject #2, shown as the example in Fig. 3). Columns show flicker frequencies of 0.5, 1, 2, 4, 8, 12, 16 and 24 Hz. Each heatmap shows the second convolutional-layer output of the two-layer 1D-CNN, consisting of 64 model-derived feature channels. Blue and black bars indicate the on/off periods of optogenetic light stimulation and natural visual stimulation, respectively. Scale bar, 1,000 ms.

**Figure S7.**
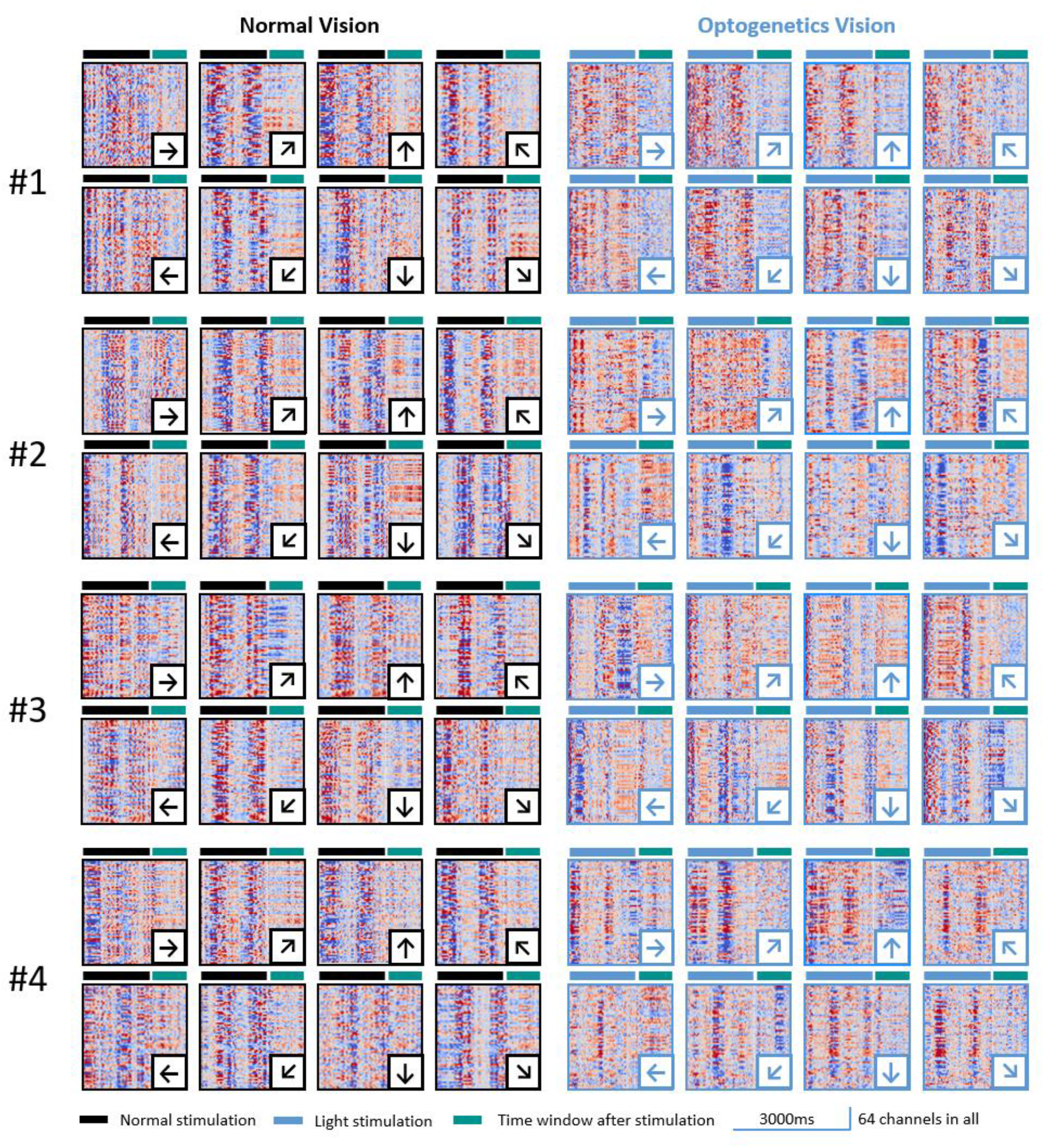
CNN feature maps of V1 responses to eight motion directions across remaining subjects. Second-layer 1D-CNN feature maps from V1 responses to drifting gratings moving in eight directions under natural vision and optogenetic artificial vision in mice #1–#4 (subject #5 is shown as the representative example in Fig. 5). Each heatmap contains 64 model-derived feature channels. Arrows indicate motion direction. Black and blue bars indicate the on/off periods of natural visual stimulation and optogenetic light stimulation, respectively; green bars mark the analysis window. Scale bar, 2,000 ms. Opto denotes optogenetic stimulation.

## References

1 Nanduri, D. et al. Frequency and amplitude modulation have different effects on the percepts elicited by retinal stimulation. Investigative ophthalmology & visual science 53, 205–214 (2012).

2 Schmidt, E. M. et al. Feasibility of a visual prosthesis for the blind based on intracortical micro stimulation of the visual cortex. Brain 119, 507–522 (1996).

3 Chen, X., Wang, F., Fernandez, E. & Roelfsema, P. R. Shape perception via a high-channel-count neuroprosthesis in monkey visual cortex. Science 370, 1191–1196 (2020).

4 Beauchamp, M. S. et al. Dynamic stimulation of visual cortex produces form vision in sighted and blind humans. Cell 181, 774–783. e775 (2020).

5 Humayun, M. S. et al. Interim results from the international trial of Second Sight’s visual prosthesis. Ophthalmology 119, 779–788 (2012).

6 Fernández, E. et al. Visual percepts evoked with an intracortical 96-channel microelectrode array inserted in human occipital cortex. The Journal of clinical investigation 131 (2021).

7 Cehajic-Kapetanovic, J., Singh, M. S., Zrenner, E. & MacLaren, R. E. Bioengineering strategies for restoring vision. Nature biomedical engineering 7, 387–404 (2023).

8 Stingl, K. et al. Interim results of a multicenter trial with the new electronic subretinal implant alpha AMS in 15 patients blind from inherited retinal degenerations. Frontiers in neuroscience 11, 445 (2017).

9 Beyeler, M. et al. A model of ganglion axon pathways accounts for percepts elicited by retinal implants. Scientific reports 9, 9199 (2019).

10 Luo, Y. H.-L. & Da Cruz, L. The Argus® II retinal prosthesis system. Progress in retinal and eye research 50, 89–107 (2016).

11 Ferlauto, L. et al. Design and validation of a foldable and photovoltaic wide-field epiretinal prosthesis. Nature communications 9, 992 (2018).

12 Beyeler, M., Rokem, A., Boynton, G. M. & Fine, I. Learning to see again: biological constraints on cortical plasticity and the implications for sight restoration technologies. Journal of neural engineering 14, 051003 (2017).

13 Boyden, E. S., Zhang, F., Bamberg, E., Nagel, G. & Deisseroth, K. Millisecond-timescale, genetically targeted optical control of neural activity. Nature neuroscience 8, 1263–1268 (2005).

14 Zemelman, B. V., Lee, G. A., Ng, M. & Miesenböck, G. Selective photostimulation of genetically chARGed neurons. Neuron 33, 15–22 (2002).

15 Bi, A. et al. Ectopic expression of a microbial-type rhodopsin restores visual responses in mice with photoreceptor degeneration. Neuron 50, 23–33 (2006).

16 Siegle, J. H. et al. Survey of spiking in the mouse visual system reveals functional hierarchy. Nature 592, 86–92 (2021).

17 Yang, R. et al. Assessment of visual function in blind mice and monkeys with subretinally implanted nanowire arrays as artificial photoreceptors. Nature biomedical engineering 8, 1018–1039 (2024).

18 Montazeri, L., El Zarif, N., Trenholm, S. & Sawan, M. Optogenetic stimulation for restoring vision to patients suffering from retinal degenerative diseases: current strategies and future directions. IEEE transactions on biomedical circuits and systems 13, 1792–1807 (2019).

19 Soltan, A. et al. A head mounted device stimulator for optogenetic retinal prosthesis. Journal of neural engineering 15, 065002 (2018).

20 Maheswaranathan, N. et al. Interpreting the retinal neural code for natural scenes: From computations to neurons. Neuron 111, 2742–2755. e2744 (2023).

21 Wang, C., Fang, C., Zou, Y., Yang, J. & Sawan, M. Artificial intelligence techniques for retinal prostheses: a comprehensive review and future direction. Journal of neural engineering 20, 011003 (2023).

22 Kriegeskorte, N. Deep neural networks: a new framework for modeling biological vision and brain information processing. Annual review of vision science 1, 417–446 (2015).

23 Zheng, Y., Jia, S., Yu, Z., Liu, J. K. & Huang, T. Unraveling neural coding of dynamic natural visual scenes via convolutional recurrent neural networks. Patterns 2 (2021).

24 Bashivan, P., Kar, K. & DiCarlo, J. J. Neural population control via deep image synthesis. Science 364, eaav9436 (2019).

25 Kubilius, J. et al. Brain-like object recognition with high-performing shallow recurrent ANNs. Advances in neural information processing systems 32 (2019).

26 Berry, M. J., Warland, D. K. & Meister, M. The structure and precision of retinal spike trains. Proceedings of the National Academy of Sciences 94, 5411–5416 (1997).

27 Cunningham, J. P. & Yu, B. M. Dimensionality reduction for large-scale neural recordings. Nature neuroscience 17, 1500–1509 (2014).

28 Van der Maaten, L. & Hinton, G. Visualizing data using t-SNE. Journal of machine learning research 9 (2008).

29 Lien, A. D. & Scanziani, M. Cortical direction selectivity emerges at convergence of thalamic synapses. Nature 558, 80–86 (2018).

30 Kobak, D. et al. Demixed principal component analysis of neural population data. elife 5, e10989 (2016).

31 Baden, T., Euler, T. & Berens, P. Understanding the retinal basis of vision across species. Nature Reviews Neuroscience 21, 5–20 (2020).

32 Foerster, O. Beitrage zur Pathophysiologie der Sehbahn und der Sehsphare. J. Psychol. Neurol., Lpz. 39, 463 (1929).

33 Penfield, W. The cerebral cortex in man: I. The cerebral cortex and consciousness. Archives of Neurology & Psychiatry 40, 417–442 (1938).

34 Brindley, G. S. & Lewin, W. S. The sensations produced by electrical stimulation of the visual cortex. The Journal of physiology 196, 479–493 (1968).

35 Dobelle, W. & Mladejovsky, M. Phosphenes produced by electrical stimulation of human occipital cortex, and their application to the development of a prosthesis for the blind. The Journal of physiology 243, 553–576 (1974).

36 Dobelle, W. H. in American Society for Artificial Internal Organs (ASAIO) Platinum 70th Anniversary Special Edition 26–49 (CRC Press, 2024).

37 Normann, R. A., Maynard, E. M., Rousche, P. J. & Warren, D. J. A neural interface for a cortical vision prosthesis. Vision research 39, 2577–2587 (1999).

38 Bosking, W. H. et al. Saturation in phosphene size with increasing current levels delivered to human visual cortex. The Journal of Neuroscience 37, 7188–7197 (2017).

39 Roelfsema, P. R., Denys, D. & Klink, P. C. Mind reading and writing: the future of neurotechnology. Trends in cognitive sciences 22, 598–610 (2018).

40 Fernández, E. & Normann, R. A. in Artificial vision: A clinical guide 191–201 (Springer, 2016).

41 Niketeghad, S. & Pouratian, N. Brain Machine Interfaces for Vision Restoration: The Current State of Cortical Visual Prosthetics: S. Niketeghad, N. Pouratian. Neurotherapeutics 16, 134–143 (2019).

42 Dong, D. W. & Atick, J. J. Statistics of natural time-varying images. Network: computation in neural systems 6, 345 (1995).

43 Palanker, D., Le Mer, Y., Mohand-Said, S., Muqit, M. & Sahel, J. A. Photovoltaic restoration of central vision in atrophic age-related macular degeneration. Ophthalmology 127, 1097–1104 (2020).

44 Stingl, K. et al. Subretinal visual implant alpha IMS–clinical trial interim report. Vision research 111, 149–160 (2015).

45 Ho, E. et al. Characteristics of prosthetic vision in rats with subretinal flat and pillar electrode arrays. Journal of neural engineering 16, 066027 (2019).

46 Granley, J. & Beyeler, M. in 2021 43rd Annual International Conference of the IEEE Engineering in Medicine & Biology Society (EMBC). 4477–4481 (IEEE).

47 Erickson-Davis, C. & Korzybska, H. What do blind people “see” with retinal prostheses? Observations and qualitative reports of epiretinal implant users. PloS one 16, e0229189 (2021).

48 Fernandez, E. Development of visual Neuroprostheses: trends and challenges. Bioelectronic medicine 4, 12 (2018).

49 Hawken, M., Shapley, R. M. & Grosof, D. Temporal-frequency selectivity in monkey visual cortex. Visual neuroscience 13, 477–492 (1996).

50 Sekirnjak, C. et al. High-resolution electrical stimulation of primate retina for epiretinal implant design. The Journal of neuroscience 28, 4446–4456 (2008).

51 Desimone, R. & Duncan, J. Neural mechanisms of selective visual attention. Annual review of neuroscience 18, 193–222 (1995).

52 Reynolds, J. H. & Heeger, D. J. The normalization model of attention. Neuron 61, 168–185 (2009).

53 Itti, L. & Koch, C. Computational modelling of visual attention. Nature reviews neuroscience 2, 194–203 (2001).

54 Atick, J. J. & Redlich, A. N. What does the retina know about natural scenes? Neural computation 4, 196–210 (1992).

55 Briggman, K. L., Helmstaedter, M. & Denk, W. Wiring specificity in the direction-selectivity circuit of the retina. Nature 471, 183–188 (2011).

56 Vaney, D. I., Sivyer, B. & Taylor, W. R. Direction selectivity in the retina: symmetry and asymmetry in structure and function. Nature Reviews Neuroscience 13, 194–208 (2012).

57 Sadtler, P. T. et al. Neural constraints on learning. Nature 512, 423–426 (2014).

58 Golub, M. D. et al. Learning by neural reassociation. Nature neuroscience 21, 607–616 (2018).

59 Gilbert, C. D. & Li, W. Adult visual cortical plasticity. Neuron 75, 250–264 (2012).

60 Shanechi, M. M. Brain–machine interfaces from motor to mood. Nature neuroscience 22, 1554–1564 (2019).

61 Silva, A. B. et al. Laminar organization of cellular microcircuits modulating human interictal epileptiform discharges. Nature Neuroscience, 1–14 (2026).

62 Geng, C. et al. Noradrenergic inputs from the locus coeruleus to anterior piriform cortex and the olfactory bulb modulate olfactory outputs. Nature communications 16, 260 (2025).

63 Klauke, S. et al. Stimulation with a wireless intraocular epiretinal implant elicits visual percepts in blind humans. Investigative ophthalmology & visual science 52, 449–455 (2011).

64 Chen, T. & Guestrin, C. in Proceedings of the 22nd acm sigkdd international conference on knowledge discovery and data mining. 785–794.

65 Pedregosa, F. et al. Scikit-learn: Machine learning in Python. the Journal of machine Learning research 12, 2825–2830 (2011).

